# Promiscuous and unbiased recombination underlies the sequence-discrete species of the SAR11 lineage in the deep ocean

**DOI:** 10.1101/2024.10.30.621061

**Authors:** Jianshu Zhao, Maria Pachiadaki, Roth E. Conrad, Janet K. Hatt, Laura A. Bristow, Luis M. Rodriguez-R, Ramon Rossello-Mora, Frank J. Stewart, Konstantinos T. Konstantinidis

## Abstract

Surveys of microbial communities (metagenomics) or isolate genomes have revealed sequence-discrete species. That is, members of the same species usually show >95% Average Nucleotide Identity (ANI) of shared genes among themselves vs. <83% ANI to members of other species while genome pairs showing between 83-95% ANI are comparatively rare. In these surveys, aquatic bacteria of the ubiquitous SAR11 clade (Class *Alphaproteobacteria*), which play a major role in carbon cycling, are an outlier and often do not exhibit discrete species boundaries, suggesting the potential for alternate modes of genetic differentiation. To explore evolution in SAR11, we analyzed high-quality, single-cell amplified genomes (SAGs) and companion metagenomes from an oxygen minimum zone (OMZ) in the Eastern Tropical Pacific Ocean, where the SAR11 make up ∼20% of the total microbial community. Our results show that SAR11 do form several sequence-discrete species, but their ANI range of discreteness is shifted to lower identities between 86-91%, with intra-species ANI ranging between 91-100%. Measuring recent gene exchange among these genomes based on a newly developed methodology revealed higher frequency of homologous recombination within compared to between species that affects sequence evolution at least twice as much as diversifying point mutation across the genome. Recombination in SAR-11 appears to be more promiscuous compared to other prokaryotic species and has facilitated the spreading of adaptive mutations within the species, further promoting the high intra-species diversity observed. These results implicate rampant, genome-wide homologous recombination as the mechanism that underlies the evolution of SAR11 into discrete species.

**Significance:** Distinguishing “species” is a pressing issue in microbiology, partly because the mechanisms that create and maintain clusters of genetically similar microbes are sparsely documented for natural populations. By leveraging high quality single-cell genomic data and a novel method for assessing homologous recombination, we show that rampant homologous recombination maintains species-level clusters of genomes for the most abundant order of marine bacteria, suggesting that these genomes may be evolving sexually to a much greater extent than previously thought. Therefore, our results identify a mechanism explaining the evolution of species in a major microbial group and have implications for understanding microbial diversity and the species concept more broadly.

## Introduction

The vast majority of Earth’s genetic diversity lies in the microbial realm. Describing how this diversity is organized is essential for identifying mechanisms of microbial evolution and predicting the functional consequences of this diversity. Genomic and metagenomic analyses of both engineered (e.g., bioremediation, wastewater treatment reactors) and natural (e.g., terrestrial or marine) systems have shown that microbial diversity is predominantly organized into “species”, representing tractable clusters of related genomes [reviewed in (1, 2)]. Typically, members of such species show ∼96-100% genome-average nucleotide identity of shared genes (ANI) among themselves and are discrete because they show <83% ANI when compared to members of other co-occurring species in the same community (3-5). More recently, our team observed other discontinuities (or gaps) in ANI values that can be used to define units within a species, most notably genomovars and strains (6, 7). Specifically, analysis of 330 diverse bacterial species each with at least 10 sequenced representative isolates revealed a scarcity of genome pairs showing 99.2-99.8% ANI (midpoint at 99.5% ANI) in contrast to genome pairs showing ANI >99.8% or <99.2%, which we suggested to refer to as genomovars (6). These analyses highlight multiple tiers of genetic organization in microbes, but do not identify the mechanisms that create and sustain this organization.

Several competing hypotheses have been advanced to explain the 95% ANI species or the intra-species gaps in microbial diversification described above. These include the hypotheses that discrete clusters are maintained by frequent recombination among closely related genomes (*recombinogenic species*) or represent differentiation into separate functional niches (*ecological species*), or combination of these two mechanisms [reviewed in (2, 8, 9)]. Recombination for prokaryotes differs fundamentally from sexual reproduction in eukaryotes in that gene exchange or shuffling does not occur during a meiosis step but via vectors of horizontal gene transfer followed by recombination of donor DNA into the recipient genome. In eukaryotes, sexual recombination underlies the “biological species concept” resulting in species cohesion. Similar genetic cohesion may also arise in prokaryotes due to recombination, resulting in what we refer to below as “recombinogenic species”. Only homologous recombination involving the replacement of an existing gene/allele by a similar foreign gene is predicted to drive unit cohesion, although homologous recombination could also drive diversification if the recombining partners represent different genomic clusters. Non-homologous recombination brings new genes and potentially new functions into the genome and leads to diversification, not cohesion. Also note that it is generally very challenging to define or measure the ecological niche of a microbial taxon, especially in natural settings, and thus directly test for the role of ecological speciation. Accordingly, rejecting recombination as the cohesive force would be sufficient to qualify shared ecology as the presumed mechanism of cohesion. Therefore, we opted to assess the level of recent recombination among the genomes reported here as a force of species cohesion (or lack of cohesion when recombination is low and/or a force of diversification) using an advanced bioinformatic approach that we recently developed for this purpose (10) as well as approaches developed by others earlier [e.g., (11)].

Comparative genomic analysis of major taxonomic groups can be vital for identifying mechanisms of diversification in microbes. The SAR11 clade of the *Alphaproteobacteria* class is one of the most ecologically dominant groups on the planet, representing up to half of the total microbial community in the surface or deep ocean (12, 13). Although the evolutionary origin of the SAR11 cluster remains uncertain (14), phylogenetic analysis of primarily the 16S rRNA gene has indicated that SAR11 bacteria form discrete (sub)clades within the broader SAR11 clade, with these subclades designated by alphanumeric identifiers (15, 16). Within these well-resolved 16S rRNA gene-based subclades, however, delineating species at the 95% ANI or another threshold has been challenging. This is primarily because mapping metagenomic reads to reference SAR11 isolate genomes or metagenome-assembled genomes (MAGs) has revealed higher intra-species genome diversity in natural SAR11 populations compared to most other prokaryotic taxa (e.g., ANI values ranging between 90-100% ANI) and sometimes indiscrete clusters or species [(3, 17, 18); and Figs. S1 and S2]. The SAR11 subclades have therefore been an outlier in seeming to lack clear species boundaries, and deviating from the 95% ANI species threshold that commonly works for the great majority of prokaryotic species (6). These observations raise the possibility of distinct modes of diversification in this major bacterial clade.

Multiple factors have presented a challenge to studying diversification in SAR11. First, reports of indiscrete SAR11 species-level diversity were based on short reads that show larger dispersion of identity values around the mean value (e.g., ANI). Second, few complete or draft genomes exist for SAR11 isolates due to challenges in isolating these bacteria, especially from deep-sea environments (19). Third, MAGs for SAR11 are typically highly fragmented and incomplete, presumably due to the higher intra- and inter-species diversity that is problematic for genome assembly and binning (20). Finally, while single-cell amplified genomes (SAGs) can provide valuable genomic data for studying SAR11 diversity, the number of available SAGs from the same or closely related SAR11 subclades is limited (18). This is again due to a high number of co-occurring species and strains in most samples, and thus not a single subclade has been sampled adequately with typical SAG efforts. That is, a single sample can contain multiple subclades, each represented by several distinct species if based on the conventional 95% ANI standard (20). Available SAG sequences are also rather incomplete (e.g., <50% completeness) and may suffer contamination issues from co-occurring cells or DNA attached to cells during cell sorting (18, 21). Therefore, a robust view of SAR11 species-level diversity based on genomes has been elusive to date, especially in sparsely sampled environments including low oxygen regions.

Studies of SAR11 from the meso- and bathypelagic realms have expanded our understanding of SAR11 diversity and its role in ocean biogeochemistry. All isolated strains of marine SAR11, including members of the ubiquitous *Pelagibacter* genus, are aerobic heterotrophs from surface or near surface waters, adapted for scavenging dissolved organic carbon (DOC) (15, 19). Their genomes are small, typically less than 1.5 Mbp, with genomic streamlining as a potential adaptation to the oligotrophic conditions of the open ocean. However, SAR11 subclades can also occur at high abundance in the meso- and bathypelagic (20, 22, 23). While it has been hypothesized that adaptation in SAR11 does not involve large variations in gene content (15), meso- and bathypelagic subclades have functional genes absent from those in surface SAR11, enabling distinct biogeochemical contributions including potential major roles in organic sulfur cycling (24) and anaerobic pathways of the marine nitrogen cycle (23). Notably, the SAR11 adapted to the marine oxygen minimum zones (OMZs) have *nar* genes supporting respiratory nitrate reduction in the absence of oxygen. The OMZ SAR11 span at least five subclades, with genomes from four of these subclades (Ic, IIa.A, IIb, and V; Fig. 1a) largely absent from non-OMZ waters but accounting for 10-30% of the bacterial community at depths with undetectable oxygen (20, 23).

**Figure 1.**
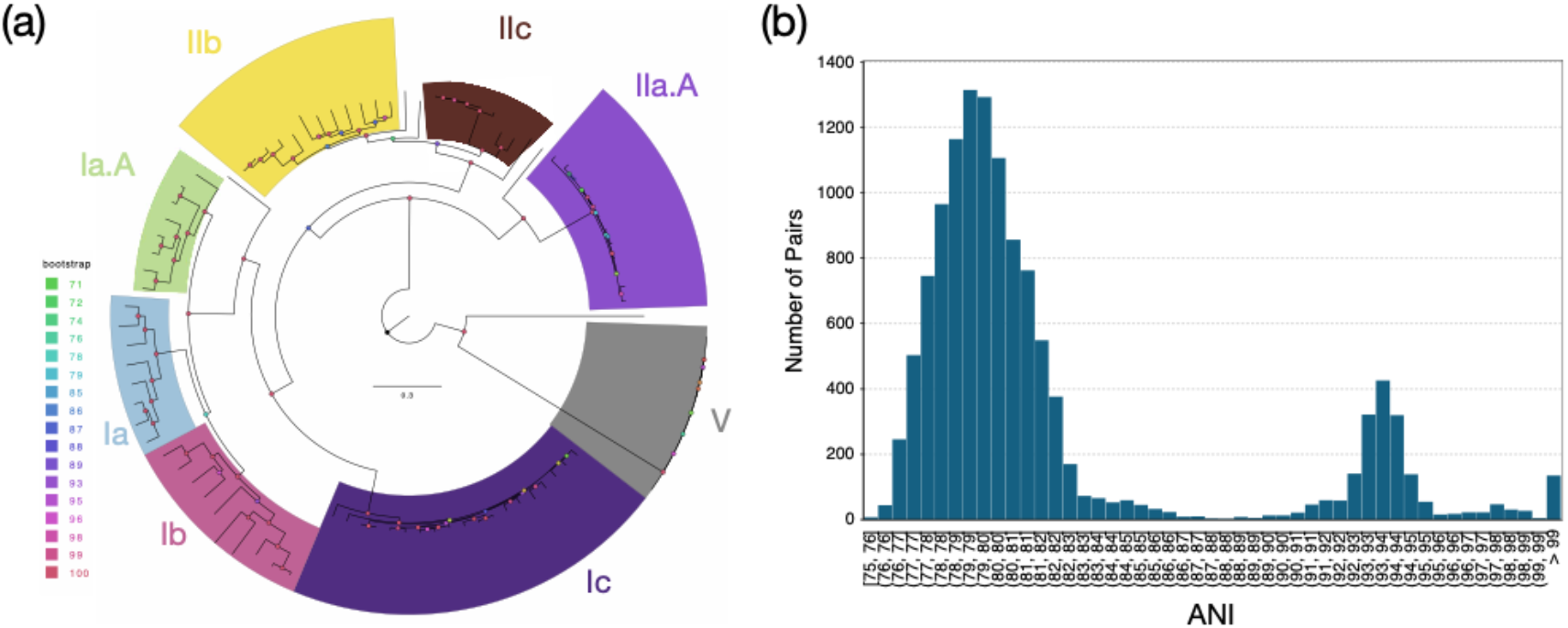
Phylogenetic and ANI diversity of the 105 SAGs used in this study. **(a)** A phylogenetic tree based on a concatenated alignment of 120 universal genes using the GToTree tool (Lee 2019). The color on the nodes of the tree indicates the bootstrap support (see legend on the left). **(b)** ANI distribution between the 105 SAGs used in the study; ANI values were calculated using FastANI (v1.3.3) (1) with default settings. Pairs of SAGs sharing less than 75% ANI are not shown because such ANI values are not reliable. Note the genomospecies gap between 86% and 91% ANI and the structure within the genomospecies (>91% ANI) with a peak in datapoints around 93-95% ANI.

The SAR11 subclades Ic and IIa are most abundant in OMZs, with the available OMZ-type genomes of subclade IIa designated as subclade IIa.A. Subclades Ic and IIa.A in OMZs show 0-3% intra-subclade differences in 16S rRNA sequences and 70-100% ANI, and thus could encompass multiple species based on the frequently used 16S rRNA gene or ANI standards (25). To provide insights into the species-level diversification process of the OMZ-abundant SAR11 subclades, we report the comparative analysis of 105, high-quality SAGs from 14 samples representing different depths along the oxycline in the Eastern Tropical North Pacific (ETNP) OMZ (see sampling details in Table S1 for SAGs and Table S5 for metagenomes). Specifically, we 1) test whether these genomes differentiate into discrete genetic clusters, 2) quantify the similarity thresholds that define cluster boundaries, and 3) and explore the forces maintaining species clusters, testing whether SAR11 clusters are maintained as cohesive units by recombination. For simplicity, we use “recombination” to refer to homologous recombination, unless noted otherwise.

## Results

### An ANI gap does exist for OMZ SAR11 species but is shifted

The average completeness and contamination of the 105 SAGs were 73.5% and 0.32%, respectively (Table S2); these values are higher than for most SAG datasets (18, 21), likely due to recent advancements in the DNA amplification protocol (26). The 105 SAGs used in this study were assigned to 8 subclades with the number assigned to each subclade roughly consistent with the relative abundance of the subclade based on metagenome read mapping (Fig. 1a, Table S2). For example, the most abundant subclade Ic in the metagenomes also contained the highest number of SAGs, suggesting that our SAG collection is a representative, random sample of the total SAR11 population *in situ*.

Pairwise comparative analysis of all SAGs revealed a species-level ANI gap (Fig. 1b), but this gap is shifted compared to the great majority of bacterial and archaeal species with adequate numbers of sequenced representatives (6) or other co-occurring species based on metagenomic read recruitment (Fig. S1b). Specifically, the ANI gap for OMZ SAR11 SAGs lies between ∼86 and ∼91%, with the intra-species ANI values ranging between 91 and 100% and exhibiting a single, prevalent peak at 93-94% (Fig. 1b). This pattern differs from the 84-96% ANI gap and 96-100% intra-species ANI range reported previously for *Escherichia coli* and other model bacterial species (6). We observed a similar ANI distribution when the analysis was restricted to the species of two abundant subclades with adequate numbers of SAGs (subclade Ic and Ib; Fig. S9). Therefore, it appears that the OMZ SAR11 community is composed of sequence-discrete species harboring high levels of genomic diversity (Fig. 1b, and Fig. S2a and b).

### Frequent, recent homologous recombination underlies the SAR11 species

We assessed the level of recent recombination among SAGs using a method recently developed in our lab (10). Briefly, our approach examines the frequency of identical genes (observed F_100_) shared between two genomes relative to the number of such genes expected by chance according to the ANI value (expected F_100_), and thus can assess if there is more recombination within vs. between clusters of genomes while normalizing for the degree of relatedness of the genomes being compared (10). By only focusing on recent exchange events (i.e., identical, or almost identical genes), our approach also circumvents computational challenges associated with historic recombination (e.g., low signal-to-noise ratio due to sequence amelioration to match the mutational biases of the recipient genome) while still being robust for assessing recombinogenic speciation. Further, for recombination to drive species cohesion it has to be frequent enough (especially compared to the effect of diversifying point mutation) and random across the genome. A non-random (biased) distribution, with recombination spatially biased to certain genomic regions or concentrated in genes of related function, could indicate selection-driven genetic exchange, mediated by recombination, but not genome introgression, since the non-recombining parts of the genome would continue to diverge.

Using this approach, we recently showed that genomes of *Salinibacter ruber*, an environmental halophilic bacterium, and *Escherichia coli*, the enteric model bacterium and opportunistic pathogen, frequently engage unbiased homologous recombination that has five times or more the effect of diversifying mutation on sequence evolution. Accordingly, recombination is responsible for maintaining species- and intra-species sequence-clusters (units) for these two species (10). Many of the SAR11 SAGs of the same species of subclade Ic (i.e., sharing >91% ANI), the most well-sampled subclade in our collection (n=30 SAGs), showed levels of homologous recombination similar or even higher than those observed previously for *E. coli* (Figs. 2 and 4). High recombination levels were observed even for SAR11 genomes that show higher divergence from each other compared to the *E. coli* genomes (ANI values ranging from 93%-94% vs. 97%-98%, respectively), despite the fact that recombination frequency is known to drop with higher sequence divergence of the recombining sequences (see also Discussion section). For instance, in pairwise comparisons among SAR11 genomes showing about 98.5% ANI, the average fraction of genes identified as recently exchanged was 30% of the total genes in the genome, whereas this percentage was 10% for *E. coli* genomes with comparable ANI (Fig. 4b) and 6% in the absence of lack of recombination (expected F_100_).

**Figure 2.**
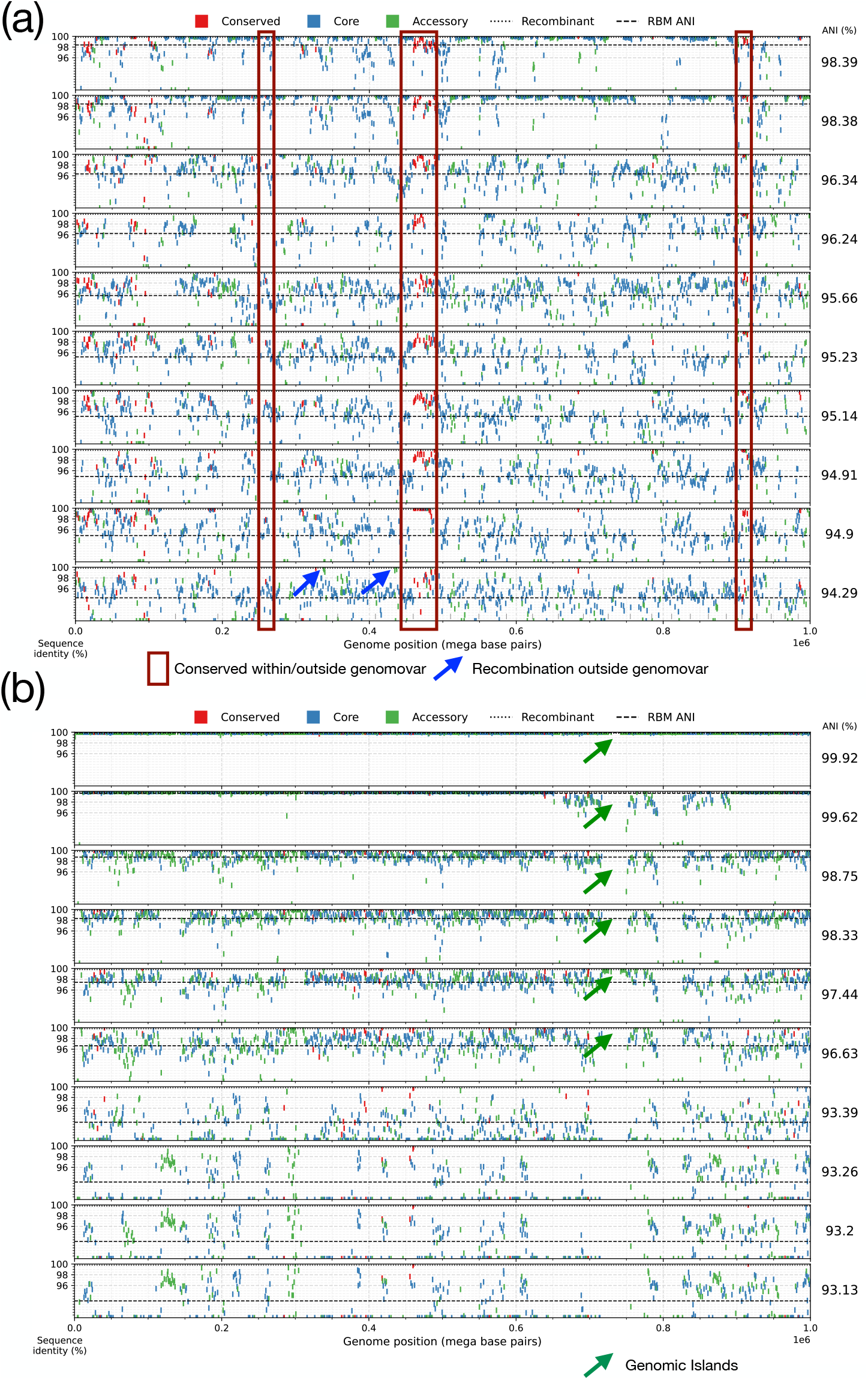
Extensive recent recombination within the SAR11 subclade Ic and comparison with *E. coli*-*E. fergusonii* genomes. (**a**) Pairwise reciprocal best match (RBM) genes were identified for 10 SAR11 subclade Ic SAGs against the same reference, SAG AM-660-D08; (**b**) similarly, for 10 *E. coli*-*E. coli* (top 6 rows) and *E. coli*-*E. fergusonii* (bottom 4 rows) genomes (reference genome is *E. coli* ASM1374039 from (2)). Each query genome is represented in a different row, and the query genomes are sorted based on decreasing ANI relatedness to the reference (rightmost values). Each rectangular marker represents a gene, colored differently for highly conserved/universal, core, and accessory genes (see figure key). The marker represents the nucleotide sequence identity between the reference and the query genome when the gene is RBM-conserved between the two genomes (y-axis) plotted against the position of the gene in the reference genome (x-axis). Blue arrows highlight genes that have most likely undergone recent recombination between the reference and the corresponding query genomes as reflected by their high nucleotide identity (>99.8%) compared to the ANI value of the genomes; green arow denote a genomic island specific to the reference genome that is not shared by most query genomes. Red boxes highlight the highly conserved *narG* and related genes discussed in the text; 1^st^ box from the left contained *narG* OP-3 type, 2^nd^ box (middle) the *narG* Gamma type. See also **Table S3** and **Table S4** for details on the genes contained within the boxes.

To further corroborate these findings, we examined the nucleotide sequence identity patterns of individual genes across the whole genome for SAGs of subclade Ic. We observed that members of the same genomovar are identical or almost identical (nucleotide identity >99.8%) in most of their genes (>50% of the total, typically), except for a few regions (hotspots) that have accumulated substantial sequence diversity (typically 90-98% nucleotide identity to other members of the same genomovar; Fig. 2, top two genomes). [Note that we used a more relaxed definition for genomovar, defined as genomes sharing >98.5% ANI, compared to our proposed definition of 99.5% ANI mentioned above and in (6) due to the lack of enough genome pairs showing >95% ANI and the higher intra-species diversity revealed for SAR11 species in Fig. 1]. Intriguingly, in almost half of the cases, genes in the diversity hotspots have an identical or almost identical match to another SAG of a different genomovar in our collection, indicating recent horizontal gene transfer (HGT) mediated by homologous recombination from that genomovar or its recent ancestors (Fig. 2, red boxes; and Fig. 5 for a phylogenetic tree-based example). It is thus likely that the other half of the genes in the diversity hotspots are also the product of recent HGT, but we did not have the donor genome within our SAG collection to confirm the HGT event (e.g., identify the high-identity match). Most of the genes in these hotspots represented core genes shared among the species, although several accessory (or variable) genes were also noted. Alternatively, these divergent genes could represent regions of hyper-mutation, but this scenario is less likely given the high identity across the rest (majority) of the genome and that the predicted functions of the divergent genes appear to be a random subsampling of the functions in the genome as a whole (Fig. 3a). Hence, the hotspots of sequence diversity between members of the same genomovar are unlikely to represent hyper-mutation or positive (adaptive) selection (see also next section for a probable case of positive selection).

**Figure 3.**
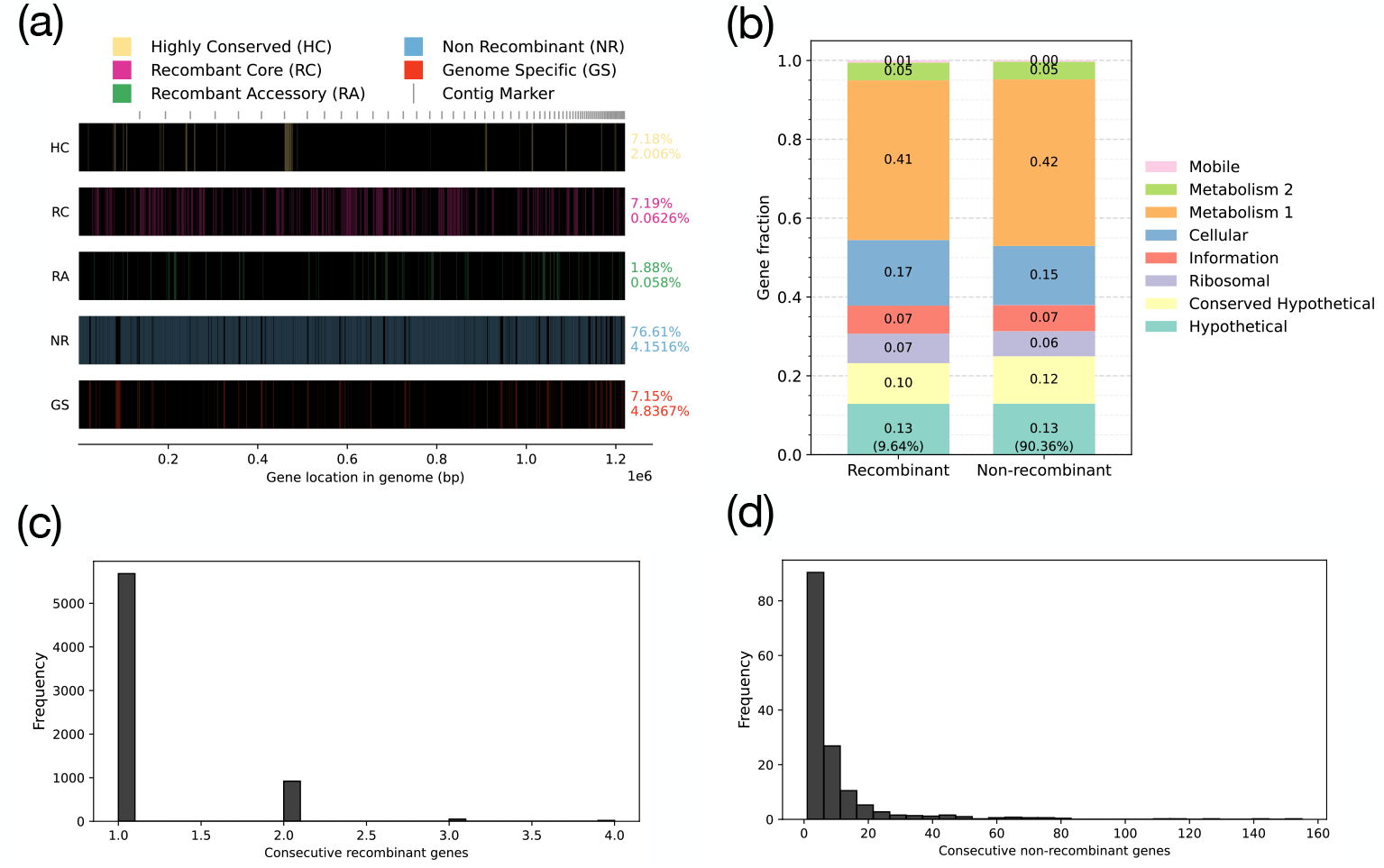
Spatial, functional and fragment length distribution analysis of recombined segments among the SAR11 SAGs. The graph shows additional analysis of the recombinant genes among the 10 genomes of the subclade Ic shown in Figure 2a. Specifically, the spatial distribution across the genome is shown in (**a**), separately for recombinant or non-recombinant genes that are core, accessary or genome-specific genes (see key on top). Functional annotation of recombinant and non-recombinant genes based on the eggNOG-mapper (version 2) is shown in (**b**). Finally, the length distribution of consecutive (**c**) recombinant (as a proxy for the length distribution of recombinant fragments) and (**d**) nonrecombinant genes are shown.

Counting all recombined genes between a reference genome and all available genomes of the subclade Ic species (i.e., cumulative recombination), not only in pairwise genome comparisons as described above, showed that SAR11 genomes have a higher cumulative recombination fraction than *E. coli* (Fig. 4). Notably, our previous study of the same *E. coli* genomes using an empirical approach to estimate the ratio of mutations purged by homologous recombination (r) vs. mutations created by point mutation over the same period of time (m), or simply the r/m ratio, revealed that the level of recombination observed above translates to an r/m ratio higher than 1 and often around 3-5 for several genome pairs (10). The SAGs analyzed here appear to have similar, if not higher, rates of recombination than the *E. coli* genomes, especially among genomes with ANI between 96% to 98%. Similar r/m ratios were also obtained with the independent approach of ClonalFrameML (Fig. S10) (11, 27). Therefore, it appears that these SAR11 SAGs are engaging in genome-wide, rapid recombination that affects sequence identity much more than diversifying (point) mutations, revealing a recombinogenic rather than clonal pattern of sequence evolution.

**Figure 4.**
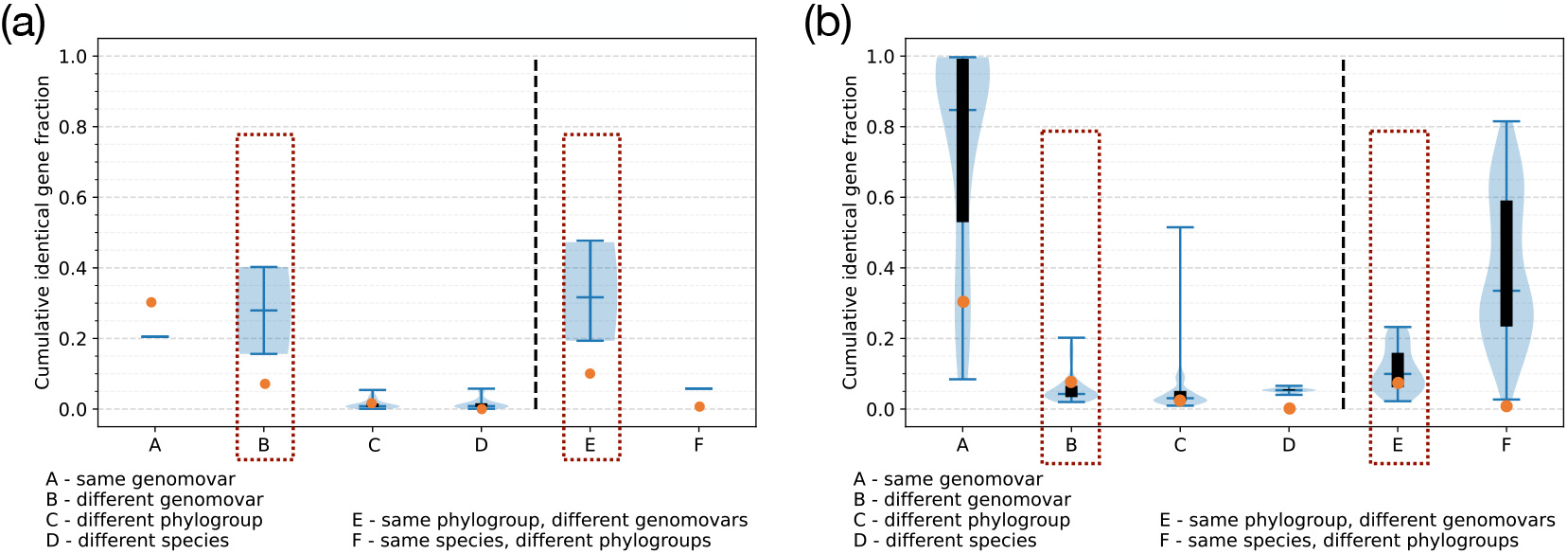
Fraction of identical genes a genome shares with all other genomes within or between genomovar, phylogroup, and species for SAR11 subclade Ic (a) and *E. coli* (b). Each genome was compared to all other genomes within each grouping (a-f) and the cumulative fraction of shared identical genes was recorded and plotted using the custom script *Allv_RBM_Violinplot*.*py*. Note that an individual gene is counted only once for this analysis so as not to overestimate recombination regardless of how many genomes were found to have recently exchanged the gene. The groupings were as follows: (a) genomes within the same genomovar defined at >98.5% ANI, (b) genomes in each separate genomovar within the same phylogroup, excluding genomes from the same genomovar, defined at 98.5%>ANI>97.5%, (c) genomes in each separate genomovar within different phylogroups defined at 97.5%>ANI>95.5%, (d) genomes of the other species (*E. fergusonii* for *E. coli*; most divergent SAGs of the same species for SAR11), defined at 95.5%>ANI>92%. Columns a through c show pairwise comparison; panels e and f show one vs many genome comparisons (cumulative). Specifically, (e) genomes within the same phylogroup excluding genomes from the same genomovar, (f) genomes within the same species excluding genomes from the same phylogroup. Data are presented in hybrid violin plots where the top and bottom whiskers show the minimum and maximum values, the middle whisker shows the median value, the black boxes show the interquartile range, and the light blue regions show the density of values along the y-axis. Note that while one or a few genomes create extreme outliers, overall, the fraction of identical genes gradually decreases among more divergent genomes compared. Also, note that our modeling analysis (orange circles on the graph; see main text for more details) suggests -for example-that only about 6-7% of the total genes in the genome should be expected to be identical among genomes showing around 98.5% ANI if there is no recent recombination (i.e., the b and e groups); both species show many more such genes in one-to-one genomovar (group b) or one-to-many genomovars (group e) at this level, revealing extensive recent gene exchange. For group a and f, there was only 1 data point for SAR subclade Ic due to lack of closely related genomes; hence, these results may not be robust but are shown for the sake of completeness and comparison to the *E. coli* data.

Substantial recent recombination was not only observed among highly related SAR11 genomes (e.g., showing >96% ANI) but also among genomes related at 93-94% ANI, albeit the measured recombination frequency was comparatively lower for the more divergent genomes (Figs. 2 and 4). It was not possible to compare these results to values for *E. coli*, given that all genomes classified as *E. coli* share > 95% ANI. However, *Escherichia fergusonii* shares ∼91-92% ANI with *E. coli*, a level comparable to that observed among the most divergent SAR11 SAGs of the same species. The *E. fergusonii* genomes analyzed here originated from the same geographic site in the United Kingdom (100 km radius) as the *E. coli* genomes (28), and thus could have engaged in more genetic exchange in comparison to genomes drawn randomly from the public databases. Nonetheless, comparing *E. fergusonii to E. coli* revealed a few hotspots of recombination, but these appear to be located in the exact same regions (genomic islands) in the available *E. fergusonii* genomes (Fig. 2b, bottom rows). Therefore, these recombination events most likely reflect selection-driven recombination or are restricted to genomic islands and are unlikely to lead to genome introgression. In contrast, recombination in the SAR11 genomes sharing 93-94% ANI occurs in many genes distributed in different regions for different query genomes, suggesting unbiased recombination (Fig. 2a, bottom rows; note the multiple high-identity genes found across the genome for SAR11 genomes but not for the *E. coli* vs. *E. fergusonii* comparison).

Finally, the length of the presumed recombined segments, using the total length of consecutive recombined genes as a proxy, was similar to that observed in previous laboratory recombination studies (29) and ranged between 1 and 20 kbp, with the majority being 1-3 kbp (Fig. 3c). Our spatial analysis also showed that, for every region of the genome longer than 100-200 kbp, the importance of recombination was greater than point mutation in at least a couple pairs of genomes from different genomovars, and there were no regions free of recombination (Fig. 3a). Further, while the fraction of the genome affected by recombination between any two genomes (of different genomovars) was almost always less than 40% of total genome length, when we compared one reference genome against representative genomes of all available genomovars and summed all recombination events (with all possible partners in the analysis), the genomic fraction affected by recombination often approached 50% or higher (Fig. 4; one vs. many comparisons). Such results were obtained with all reference genomes and were not specific to any subclade. Therefore, it appears that for the SAR11 subclade Ic species evaluated here, homologous recombination is frequent and random enough (spatially across the genome) and likely sufficient to serve as the mechanism for species cohesiveness. While these results are predominantly based on subclade Ic for which we had the highest number of SAGs, similar frequency of recombined genes and a lack of spatial bias across the genome were observed for SAGs representing the surface (non-OMZ) (Fig. S5), suggesting these patterns may be broadly applicable across SAR11 species from distinct oceanic niches.

### Gene sweeps, rather than genome sweeps, shape SAR11 functional diversity, consistent with recombinogenic species and high intra-species diversity: the case of the nitrate reductase (narG)

While recombination affected almost every gene in the genome (Fig. 2a), we also observed three hotspots where recombination was more frequent than in the rest of the genome and consequently resulted in low sequence divergence of the affected genes (Fig. 2a). Two of the hot spots encompassed genes encoding the two variants (OP3 and Gamma variants) of respiratory nitrate reductases (NarG) previously identified as distinguishing OMZ SAR11 subclades from their counterparts in oxic water columns (23). The 3^rd^ hotspot included genes of other metabolic functions (Table S4). The first hotspot encompassed only the *narG* gene encoding the OP3 nitrate reductase variant, while the second hotspot included the entire operon (*narGHIJ;* Table S3) encoding the Gamma Nar variant. The OP3 and Gamma *narG* variants share only about ∼45% amino acid identity (no detectable identity or alignment at the nucleotide level), and therefore are unlikely to undergo homologous recombination with each other. Notably, genes in the two *nar*-encoding hotspots showed very high sequence identity across SAGs, ranging from ∼99% to 100%, even among SAGs that show 93-94% total genome ANI. A plot showing metagenomic reads recruited to the contig containing the *narGHIJ* operon confirmed that the operon exhibits consistent levels of high similarity across the *in situ* SAR11 community (Fig. S7), not only among our SAGs.

This level of high sequence identity (>99%) in *nar* genes is comparable to only that for rRNA genes, which are known to be highly conserved due to functional constrains (Fig. 5 and Table S3). Such high similarity is extremely rare for functional genes such as *nar* (30), unless the genes are recently horizontally exchanged or the genomes are very closely related (e.g., >99% ANI; thus, not enough evolutionary time for sequence diversification), which is not the case for most SAR11 SAGs analyzed here. The phylogenies of both *narG* variants were not congruent with that based on corresponding 16S rRNA gene sequences, further suggesting HGT. In fact, when we compared the *narG* and 16S rRNA gene phylogenies against an ANI-based or a core-genome tree, we observed at least two cases of clear HGT of not only *narG* but also the 16S rRNA gene within (but not outside) subclade Ic, assuming the ANI tree best represents the species phylogeny (Fig. 5). These results are consistent with the rampant and unbiased recombination described above, notably as rRNA genes are transferred horizontally only rarely (31). Collectively, these results suggest that *narG*, and likely the entire *nar* operon, has undergone a gene sweep event within the subclade Ic species, presumably caused by positive (adaptive) selection for the prevailing *narG* allele (allele is defined here as sequences of the gene showing >98.5% nucleotide sequence identity, Table S3). While frequent recombination has seemingly kept *nar* sequence diversity low across the genomes, the apparent selective advantage of *nar* has not caused sweep events at the level of the genome - genomes with *nar* did not outcompete and drive extinct genomes without *nar*. If a genome sweep event had taken place, the remainder of the genome would be expected to show low sequence divergence similar to that of the *nar* operon (i.e., >99% ANI vs. 91-100% ANI observed, with the median around 94%). Previous research has shown nitrate reduction, mediated by Nar, to be a key energy-generating pathway for OMZ-associated SAR11 across all OMZ SAR11 subclades (23), consistent with the presumed strong positive selection for Nar function.

**Figure 5.**
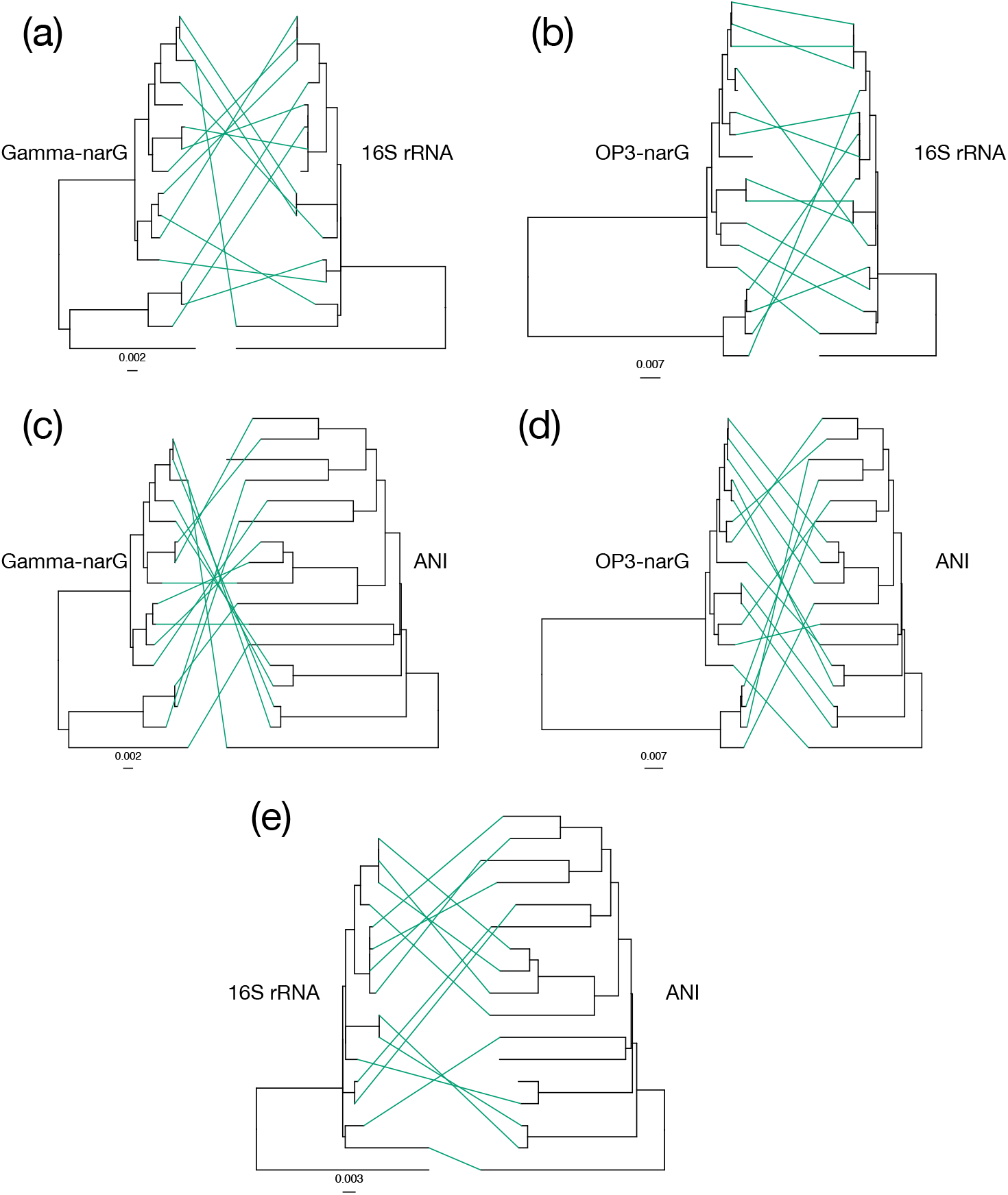
Phylogenetic relationships among subclade Ic SAGs based on ANI, 16S rRNA and *narG* genes. See labels on top of each tree for which gene (or genome ANI) the tree represents. In short, the top row represents (**a**) Gamma-type *narG* (second hot spot in Figure 2a) and (**b**) OP3-type *narG* (first hot spot in Figure 2a) in comparison to the 16S rRNA gene, whereas the second row (**c** and **d**) shows the same two *narG* gene trees as in a and b in comparison to an ANI tree. The last row shows the 16S rRNA gene in comparison to the ANI tree. The scale bars denote the substitution rate. All trees are built using nucleotide sequences and the neighbor-joining algorithm. The long, unpaired branch in 16S rRNA and *narG* trees represents subclade V and was used as the outgroup. All trees are mid-point rooted. Note the similar level of sequence diversity within the *narG* and the 16S rRNA genes, and that several cases of probable horizontal gene transfer of these genes can be observed when taking the ANI as the species tree (e.g., green lines representing different topologies compared to the ANI tree).

## Discussion

The SAG sequences analyzed here corroborate previous results (3, 20) indicating that OMZ SAR11 populations are organized in sequence-discrete species but with higher intra-species diversity than most other bacteria studied to date, i.e., ANI values in the range of 91-100% vs. 96-100% for most studied species (6). Further, the analysis revealed that homologous recombination may be the force driving species cohesion for these genomes as recombination was found to be both frequent (i.e., greater impact on sequence evolution compared to diversifying mutation) and random across the genome. In contrast, non-homologous recombination that could bring new functions into the genome was at least 10 times less frequent among the SAGs (Fig. 2), suggesting that ecological speciation is less important for species cohesion compared to recombinogenic speciation. Our results also explain several of the challenges reported in the recent literature for those studying species-level diversity in SAR11. That is, the high level of intra-species sequence diversity in SAR11 (Fig. 1b) is outside the range of most assembly programs that typically merge sequences that share at least 97-98% identity, which presumably accounts -in part-for the highly incomplete SAR11 MAGs recovered previously (20). Further, the pattern of indiscrete species revealed previously is largely due to artifacts of the short-read sequence analysis. For example, short reads show larger dispersion around the mean nucleotide identity value (e.g., ANI) when SNPs accumulate over relatively short pieces of DNA, which, given the level of sequence diversity revealed among our SAGs, is rather common within SAR11 species.

Previous experimental studies have shown that the frequency of recombination drops substantially around 95% nucleotide identity albeit recombination is still possible even among pieces of DNA that are less than 90% identical, depending on the secondary structure of the recombining DNA and the promiscuity of the corresponding enzyme systems, among other factors (32). More recently, high-throughput transformation studies at the whole-genome level have further corroborated these earlier findings and showed that homologous recombination efficiency in *Bacillus subtilis* drops by 5-fold or more for sequences that share 94-95% nucleotide identity relative to those that share 99-100% identity (29). This prior work has also shown that homologous recombination efficiency drops to almost zero (∼20-fold compared to 99-100% identical sequences) around 88-90% identity, which intriguingly corresponds to the highest intra-species divergence level observed for the SAR11 genomes studied here (91-100% ANI; if the ANI is 91%, this would mean that several individual genes in the genome show 88-90% nucleotide identity). There are two additional lines of evidence that further corroborate that SAR11 bacteria are able to recombine even sequences that are around 90% nucleotide identity. First, all genomes have a least a few protein-coding genes with a 99-100% nucleotide identity match with a second genome that shares around 94% ANI in the rest of the (shared) genes (i.e., a genome representing a divergent genome, but of the same species), which represents direct evidence of recent exchange of these genes with the latter genome (Fig. 2). Second, when we examined the underlying identity distribution of shared genes between two genomes showing high ANI (98.5%), we detected several genes with lower nucleotide identities of ∼90-92%. Such genes made up a larger fraction of the total genes in the genome in SAR11 when compared to *E. coli* genomes of similar ANI, causing the distribution to deviate further from a normal distribution around the mean value (the ANI value, Fig. S6). It is unlikely that these represent genes with an increased rate of fixed mutations (increased mutation rate) for the reasons mentioned above but also because several of these genes encode core functions. Rather, a more parsimonious scenario is that such genes are the products of recombination involving the most divergent members of the species. In conclusion, it appears that SAR11 bacteria do form sequence-discrete species, and that homologous recombination is the major force of species cohesion, even though several members of the species show sequence relatedness near the lowest limit at which homologous recombination is still possible (i.e., around 90% nucleotide identity).

It is intriguing that SAR11 shows higher intra-species diversity compared to other bacterial species. This could be due to a relatively high amount of evolutionary time since the last genome sweep event but also to higher recombination promiscuity, notably if the identity threshold for homologous recombination is comparatively low in SAR11. Consistent with the latter interpretation, OMZ SAR11 species, similar to their surface *Ca*. Pelagibacter ubique relatives, are remarkably different from the other alphaproteobacterial species in the repertoire of their genes for homologous recombination and repair. Specifically, genes encoding the RecBCD complex, which initiates homologous recombination by helping to load the primary homologous recombination protein RecA onto double-stranded gaps, are absent from both SAR11 Ic and IIa.A representatives. Similarly, among *Alphaproteobacteria, Ca*. Pelagibacter ubique and subclade Ic are unique in that they lack the RecF pathway for homologous recombination associated with DNA repair (Fig. S8), suggesting that these bacteria might utilize the recently proposed RecF-independent pathway, the RecOR pathway, for the loading of RecA for single-strand gap repair. It is hypothesized that the simplified RecOR pathway might initiate homologous recombination at both single-stranded and double-stranded breaks [reviewed in (33)], potentially resulting in a sequence identity threshold for recombination different from that of other *Alphaproteobacteria*. Further and perhaps more importantly, two additional key genes involved in the mismatch repair system - *mutS* and *mutL* - are missing from the SAGs of subclades Ic and IIa.A (Fig. S8), despite their presence in all other *Alphaproteobacteria* genomes. The MutSL protein complex corrects base pair mismatches and acts as an anti-recombinant by preventing homologous recombination of slightly diverging sequences (33). Collectively, these results indicate that SAR11 species may be evolving under enhanced rates of nucleotide substitutions and more promiscuous recombination, which could account, at least partially, for the higher diversity of ANI values observed within species (e.g. Fig. 1a and b). It would be interesting to test this hypothesis in the future using experimental studies (e.g., of DNA transformation rates).

The intra-species genetic structure observed for SAR11 might also be explained by a long period of time since the last diversity-purging selection event. Members of the species could have been evolving together since this event, accumulating point mutations but also engaging in frequent homologous recombination. The fact that most SAGs share 93-95% ANI and only a few share >96% (Fig. 1b) is consistent with this scenario of a long time since the last population/genome sweep and frequent recombination between relative divergent genomes showing 93-95% ANI. [It is also important to recognize that 1% nucleotide difference (corresponding to 99% ANI) represents thousands of generations (and likely many years) since the last common ancestor, assuming no recombination (34).] It follows that genomes that accumulate additional mutations beyond this species boundary (i.e., they diverge below the 91-92% ANI threshold) could either be driven extinct by competition with other members of the species or diverge to represent distinct species, potentially accounting for the 86-91% ANI gap (Fig. 1). The former scenario of being outcompeted could be driven by a lack of recombination with the remaining members of the species, with this lack of recombination involving advantageous alleles or genes presumably representing an important disadvantage. The results involving nitrate reduction-encoding *nar*, with all members of the 91-100% ANI species cluster containing highly identical *narG* sequences (>97.5% nucleotide identity, Table S3) and even SAGs from different subclades showing high *nar* relatedness (Fig. S4), is compatible with our interpretation in the latter scenario. Any OMZ SAR11 genome unable to acquire these *nar* alleles - likely via homologous recombination - is at a serious disadvantage under this scenario, highlighting the importance of nitrate respiration in these OMZ-adapted SAR11 lineages.

In conclusion, our results show that while the SAR11 clade is an outlier with respect to its level of intra-species genome diversity, this major bacterial group does diverge via many sequence-discrete species. The ANI threshold delimiting these species is notably lower (91-92%) than that observed for most other major bacterial lineages studied to date. The cohesive force for speciation in SAR11 appears to be homologous recombination, and the higher intra-species diversity appears due -at least in part-to the SAR11 species being more promiscuous, with recombination commonly occurring between genomes sharing relatively low identity. This promiscuity could be mechanistically attributable to a unique recombination apparatus in SAR11 compared to other *Alphaproteobacteria*, although this hypothesis would require experimental validation. Our study provides a robust, new methodology for assessing species- and intra-species-level diversity and the impact of recombination on sequence evolution in microorganisms.

## Materials and Methods

Details of all methods used in this study are described in the Supplementary Material, including how the SAGs were obtained and sequenced, how data were processed and quality-checked, and how bioinformatic analysis of the resulting data was performed. Additional references provide further information about procedures and analytical techniques.

## Data availability

Metagenomic raw data can be found in NCBI (project accession number PRJNA1124864). Raw sequence of single amplified genomes can be found in NCBI (PRJNA1124867).

## Author contributions

KTK and FS designed the work. JZ performed the bioinformatic analysis of SAG and short-read data. MP did the SAG sampling and sequencing, and assisted in their sequence analysis and interpretations of the results. REC developed the recombination detection and quantification pipeline. FS and LB did the metagenomic sampling. JKH did the metagenomic sequencing. JZ and KTK wrote the manuscript. All authors read and edited of the manuscript.

## Acknowledgement

We want to thank PACE (The Partnership for an Advanced Computing Environment) at Georgia Tech for providing computational resources, Beate Kraft and Jennifer Crandall for their help with sampling and DNA extraction, and Mark Altabet and the crew of the *R/V Sally Ride* for making sample collection possible. This work is supported by NSF (Awards 1831582 and 2129823 to KTK and 2130185 and 2022991 to FJS) and the Simons Foundation (Aquatic Microbial Ecology Award to FJS).

## Reference cited

1. A. Caro-Quintero, K. T. Konstantinidis, Bacterial species may exist, metagenomics reveal. Environ Microbiol 14, 347–355 (2012).

2. K. T. Konstantinidis, Sequence-discrete species for Prokaryotes and other microbes: A historical perspective and pending issues. mLife 10.1002/mlf2.12088, In press (2023).

3. K. T. Konstantinidis, E. F. DeLong, Genomic patterns of recombination, clonal divergence and environment in marine microbial populations. ISME J 2, 1052–1065 (2008).

4. M. R. Olm et al., Consistent Metagenome-Derived Metrics Verify and Delineate Bacterial Species Boundaries. mSystems 5 (2020).

5. M. L. Bendall et al., Genome-wide selective sweeps and gene-specific sweeps in natural bacterial populations. ISME J 10, 1589–1601 (2016).

6. L. M. Rodriguez-R et al., An ANI gap within bacterial species that advances the definitions of intra-species units. mBio 10.1128/mbio.02696-23, e0269623 (2024).

7. T. Viver et al., Towards estimating the number of strains that make up a natural bacterial population. Nat Commun 15, 544 (2024).

8. C. Fraser, W. P. Hanage, B. G. Spratt, Recombination and the nature of bacterial speciation. Science 315, 476–480 (2007).

9. B. J. Shapiro, M. F. Polz, Microbial Speciation. Cold Spring Harb Perspect Biol 7, a018143 (2015).

10. R. E. Conrad, Brink, C. E., Viver, T., Rodriguez-R, L. M., Aldeguer-Riquelme, B., Hatt, J. K., Venter, S. N., Rossello-Mora, R., Amann, R., and Konstantinidis, K. T., Microbial species and intraspecies units exist 1 and are maintained by ecological cohesiveness coupled to high homologous recombination. Nature Communications, In press. (2024).

11. X. Didelot, D. Falush, Inference of bacterial microevolution using multilocus sequence data. Genetics 175, 1251–1266 (2007).

12. R. M. Morris et al., SAR11 clade dominates ocean surface bacterioplankton communities. Nature 420, 806–810 (2002).

13. C. A. Carlson et al., Seasonal dynamics of SAR11 populations in the euphotic and mesopelagic zones of the northwestern Sargasso Sea. ISME J 3, 283–295 (2009).

14. H. Luo, Evolutionary origin of a streamlined marine bacterioplankton lineage. ISME J 9, 1423–1433 (2015).

15. J. C. Thrash et al., Single-cell enabled comparative genomics of a deep ocean SAR11 bathytype. ISME J 8, 1440–1451 (2014).

16. P. Yarza et al., Uniting the classification of cultured and uncultured bacteria and archaea using 16S rRNA gene sequences. Nat Rev Microbiol 12, 635–645 (2014).

17. T. O. Delmont et al., Single-amino acid variants reveal evolutionary processes that shape the biogeography of a global SAR11 subclade. Elife 8 (2019).

18. M. G. Pachiadaki et al., Charting the Complexity of the Marine Microbiome through Single-Cell Genomics. Cell 179, 1623–1635 e1611 (2019).

19. S. J. Giovannoni et al., Genome streamlining in a cosmopolitan oceanic bacterium. Science 309, 1242–1245 (2005).

20. C. A. Ruiz-Perez et al., Description of Candidatus Mesopelagibacter carboxydoxydans and Candidatus Anoxipelagibacter denitrificans: Nitrate-reducing SAR11 genera that dominate mesopelagic and anoxic marine zones. Syst Appl Microbiol 44, 126185 (2021).

21. T. Ishoey, T. Woyke, R. Stepanauskas, M. Novotny, R. S. Lasken, Genomic sequencing of single microbial cells from environmental samples. Curr Opin Microbiol 11, 198–204 (2008).

22. K. T. Konstantinidis, J. Braff, D. M. Karl, E. F. DeLong, Comparative metagenomic analysis of a microbial community residing at a depth of 4,000 meters at station ALOHA in the North Pacific subtropical gyre. Appl Environ Microbiol 75, 5345–5355 (2009).

23. D. Tsementzi et al., SAR11 bacteria linked to ocean anoxia and nitrogen loss. Nature 536, 179–183 (2016).

24. H. J. Tripp et al., SAR11 marine bacteria require exogenous reduced sulphur for growth. Nature 452, 741–744 (2008).

25. K. T. Konstantinidis, R. Rossello-Mora, R. Amann, Uncultivated microbes in need of their own taxonomy. ISME J 11, 2399–2406 (2017).

26. R. Stepanauskas et al., Improved genome recovery and integrated cell-size analyses of individual uncultured microbial cells and viral particles. Nature Communications 8, 84 (2017).

27. X. Didelot, D. J. Wilson, ClonalFrameML: efficient inference of recombination in whole bacterial genomes. PLoS computational biology 11, e1004041 (2015).

28. L. P. Shaw et al., Niche and local geography shape the pangenome of wastewater- and livestock-associated Enterobacteriaceae. Sci Adv 7 (2021).

29. J. J. Power et al., Adaptive evolution of hybrid bacteria by horizontal gene transfer. Proc Natl Acad Sci U S A 118 (2021).

30. K. T. Konstantinidis, J. M. Tiedje, Towards a genome-based taxonomy for prokaryotes. Journal of Bacteriology 187, 6258–6264 (2005).

31. S. R. Miller et al., Discovery of a free-living chlorophyll d-producing cyanobacterium with a hybrid proteobacterial/cyanobacterial small-subunit rRNA gene. Proc Natl Acad Sci U S A 102, 850–855 (2005).

32. M. Vulic, F. Dionisio, F. Taddei, M. Radman, Molecular keys to speciation: DNA polymorphism and the control of genetic exchange in enterobacteria. Proc Natl Acad Sci U S A 94, 9763–9767 (1997).

33. J. Viklund, T. J. Ettema, S. G. Andersson, Independent genome reduction and phylogenetic reclassification of the oceanic SAR11 clade. Mol Biol Evol 29, 599–615 (2012).

34. J. G. Lawrence, H. Ochman, Molecular archaeology of the Escherichia coli genome. Proc Natl Acad Sci U S A 95, 9413–9417 (1998).

