## Supplementary Figures and Tables for "Promiscuous and unbiased recombination underlies the sequence-discrete species of the SAR11 lineage in the deep ocean"

### Supplementary Material

#### Material and Methods

*Collection of ETNP sea water samples*

Samples for metagenomic sequencing were collected from the ETNP OMZ (R/V Sally Ride (SR2114), December 2021 – January 2022). Sea water for metagenomes was collected from all nine depths for all five sampling stations (See Table S5 for details). Collections were made using Niskin bottles on a rosette containing a conductivity–temperature–depth profiler (Sea-Bird SBE 911plus), as described previously (1).

*Water sampling and Single Cell Amplified Genome (SAG) generation*

Samples for single cell sorting and genomic sequencing were collected during the oceanographic expedition AT50-08 onboard *R/V Atlantis* at Eastern Tropical North Pacific (ENTP) Ocean. Sea water for single-cell sorting and single amplified genome (SAG) analysis was collected from two stations. Station coordinates and collection dates of each sample are described in Table S2. Water samples were prepared by cryopreservation according to the protocol recommended by the Bigelow Single Cell Genomics Center (see details below). Briefly, triplicate 1 ml samples of bulk sea water (no prefiltration) were gently mixed with 100 µl of a glycerol TE stock solution (20 ml 100× TE pH 8.0, 60 ml sterile water, 100 ml glycerol) and frozen at -80 °C. Water was sampled from the Niskin bottles immediately upon recovery, and 1ml subsamples were transferred to cryovials. For water samples from low oxygen concentrations or water samples in which oxygen was undetectable (all samples besides the one recovered from 1000m), the transfer was performed within an anaerobic glove bag. Additionally, the cryovials used had been stored in glass bottles filled with helium gas to off-gas oxygen. To select for viable cells or cell with respiratory activity, the subsamples were amended with 1µL BacLight™ RedoxSensor™ Green (Thermofisher Scientific) following the protocols of the Single Cell Genomics Facility (SCGC), Maine, USA and Lindsay, *et al.* (2). After short vortexing (1-2 secs), the vials were incubated in the dark at in situ temperatures for 30 mins. Following incubation 100ml of glyTE buffer was added (5% glycerol, 10 mM Tris, and 0.1 mM EDTA final concentration). Vials were frozen at -80˚C and shipped to the SCGC for sorting. Sorting was performed on April 4$^{th}$, 2023 (within less than two months from the date the first sample was collected) and SAGs were generated with the modified genomic DNA amplification technique, WGA-Y, which enables a substantially improved average genome recovery from single cells (service S-202). Around 100 SAGs with Cp values less than 3h were randomly selected for sequencing. Genome assembly and draft annotation were performed by SCGC as described in the center's webpage https://scgc.bigelow.org/capabilities/service-description/. Briefly, SAG paired-end libraries were created with Nextera XT kits (Illumina), sequenced with a NextSeq 500 (Illumina) and de novo assembled using a workflow utilizing SPAdes (3), as previously described (4). The total number of SAGs assembled was 1152, 128 of which were affiliated to Pelagibacterales using GSearch (v1.3.0) against GTDB database v214.

*Metagenomic sequencing and analysis*

DNA was extracted from sea water samples following a similar protocol as previously described (5). Briefly, DNA was extracted from biomass on collecting filters using the MoBio Power Soil kit (MoBio Inc. Carlsbad, CA, USA) and libraries were prepared for metagenomic sequencing using the Illumina DNA library prep kit with unique dual indexing according to manufacturer's instructions, except that the protocol was terminated after isolation of cleaned double stranded libraries. An equimolar mixture of the libraries was sequenced on an Illumina NovaSeq 6000 instrument at the Molecular Evolution Core, Georgia Institute of Technology. A negative control was used to make sure that our samples are not contaminated during the extraction process. Initial adapter trimming and demultiplexing of sequenced samples was carried out by the instrument. Adapter trimming and quality control of short reads was performed via fastp and then checked via Falco via MiGA pipeline (6). Metagenomic coverage was estimated via the Nonpareil software v3.4 (-kmer option) (7). IDBA-UD was then used to assemble short reads into contigs (--min_contig 1000) (8). Recruitment plot of short reads against MAGs and SAGs were created using the corresponding tool from the enveomics package (9), which minor modifications (see: https://github.com/jianshu93/RecruitmentPlot_blast). Metagenomic binning was then performed via the binning module of MetaWRAP, which is a wrapper that integrates three widely used binning software MaxBin2, metaBAT2 and CONCOCT (10). MAGs/bins from the binning module were then refined using DASTools (v1.1.2) (11). Quality control was performed using checkM (12) while taxonomic classification was obtained via GSearch (v1.3) (13) against the GTDB database v214 (14).

*Recombination analysis*

We developed a bioinformatic pipeline to analyze those SAGs and identify gene exchange/recombination events. Step by step details for our main analysis workflow can be found here: https://github.com/rotheconrad/F100_Prok_Recombination. Briefly, the pipeline identifies reciprocal best matches (RMBs) via BLAST (15) and then uses a metric called F100 (100% identical RBMs divided by total RBMs) as a proxy for strength of recent gene exchange events. ClonalFrameML was used as an independent method to identity recombination events and estimate recombination to mutation rate (r/m) based on coalescent models as described previously (16).

*Phylogenetic analysis and functional annotation*

SAG genome trees were built using either GToTree (based on a concatenated universal gene alignent) (17) or ANI distance matrices produced by FastANI (18), while gene trees were built via IQ-TREE2 (19) after aligning the corresponding sequences with MUSCLE (20). Genes were extracted from SAGs using protein BLAST and annotated with eggNOG-mapper (v2.1.12) (21) against the eggNOG Database (v6.0) (22).

*Data visualization*

All plots were obtained via R ggplot2 library (v3.4.4) or python Matplotlib library (v3.9.0).

### Supplementary Figures

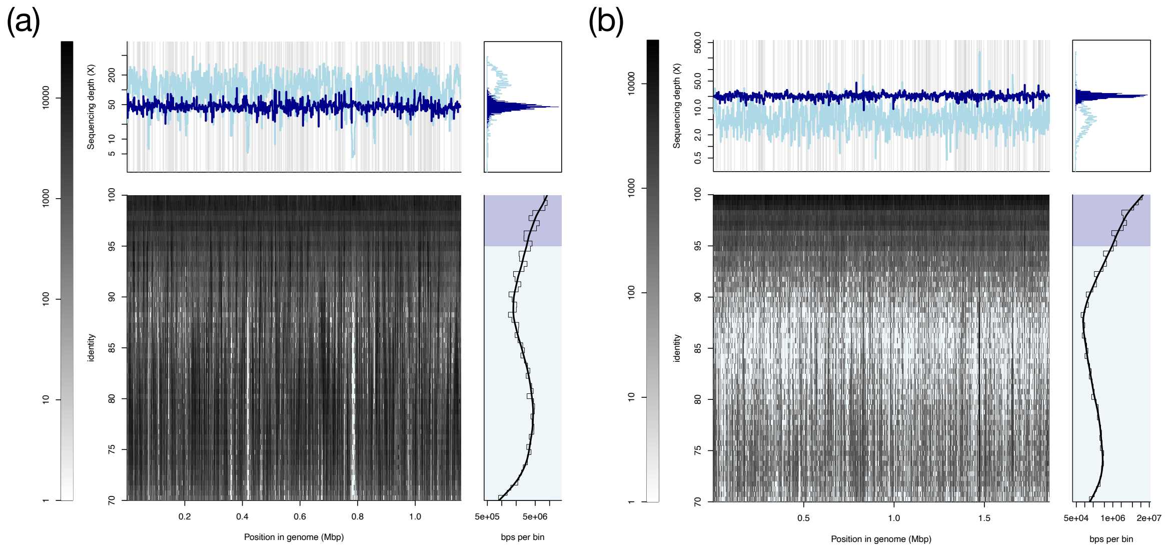


**Figure S1**. **Recruitment plot for (a) a SAR11 MAG (subclade Ic) and (b) an *Acidimicrobiales* sp. MAG against the same metagenomic sample from station 38, depth 302 m from which the MAGs originated**. The SAR11 MAG quality was estimated to be 89.5% complete and 13.2% contaminated. The main panel of (a, bottom center) shows the position (x-axis) and identity (y-axis) of the metagenomic reads that map on the reference MAGs, which are summarized into two categories: reads above 95% (dark blue) and below 95% (light blue) sequence identity. These two categories are maintained in the other panels. The lower - right panel shows total reads mapped (in base pairs, log scale) for different identities across the entire genome while the upper panels show the coverage of MAGs by reads at each position across their sequence. Note that the sequencing coverage is in log scale. Note the clear, sequence-discrete population for the *Acidimicrobiales* sp. around 80% to 95% nucleotide identity, but the much less discrete SAR11 population. See reference (23) for further details on the recruitment plots.

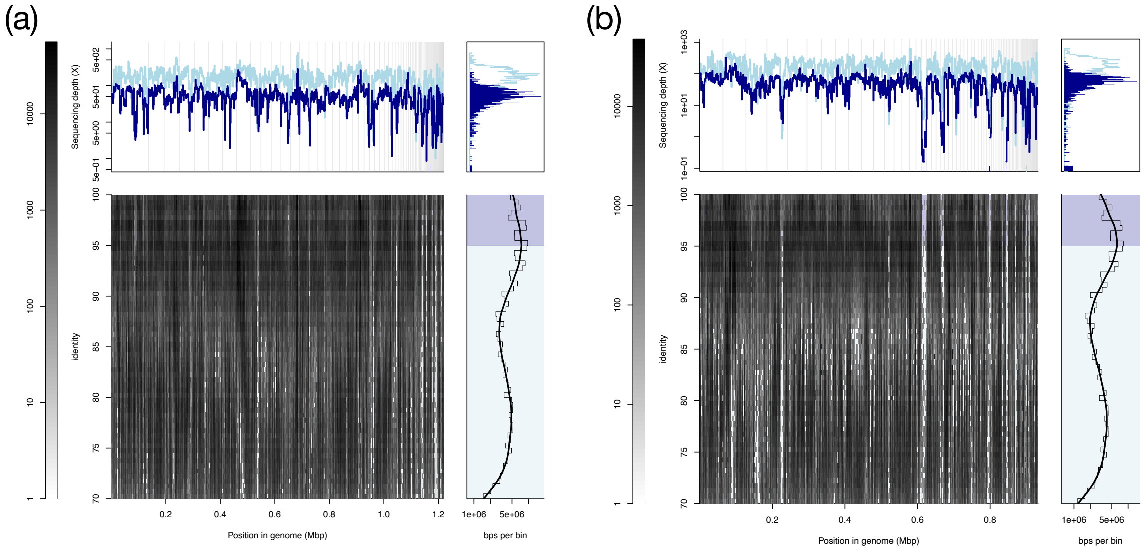


**Figure S2**. **Recruitment plot of a SAR11 SAG of subclade (a) Ic and (b) IIa.A against the same metagenomic sample from station 38, depth 302 m**. Figure S2 is identical to Figure S1 except that the best quality SAG for each subclade was chosen as a reference genome for mapping of the metagenomic reads. Note that the peak in reads mapping around 95% is consistent with the ANI distribution revealed when comparing all SAGs (i.e., Figure 1), and that the total population is rather indiscrete, especially in comparison to Figure S1(b), although a weak valley can be observed around 90% nucleotide identity (similar to Figure 1b based on whole SAG sequences).

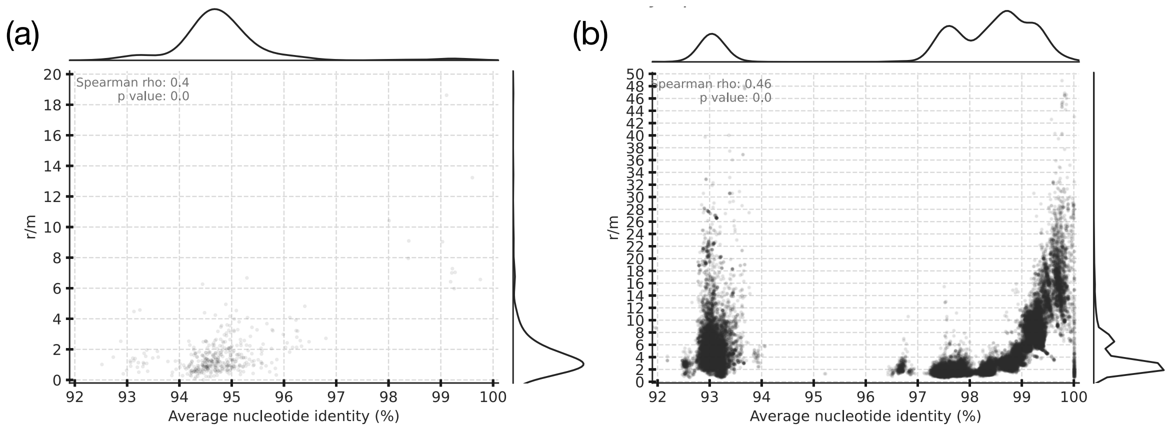


**Figure S3. Recombination to mutation ratio (r/m) versus ANI for (a) SAR11 subclade Ic and (b) *E. coli*.** See Methods & Materials for how r/m was calculated. The r/m ratio (y-axes) was estimated for all genome pairs in our collection for each species and is plotted against the ANI value of the genome pair compared (x-axes). The marginal plots outside the two axes show histograms for the density of datapoints on each axis. Note that the ratio is frequently above 1, especially for genomes sharing between 97.5-99.5% ANI (e.g., members of different genomovars of the same phylogroup) for both species and that the estimates above ~99.5% ANI are not reliable due to inability to detect recombination at this high sequence identity level. A few outlier datapoints (genome pairs) with ratios higher than 100 were also observed in the 98-99.5% ANI range and are due to the high identity of the recombined genes identified (causing the denominator in the r/m ratio to be a small number); the majority of datapoints are between ~1 and 20. Also note that a few *E. coli* and *E. fergusoni* genome pairs (left part of the graph on the right) show a ratio higher than 1, but this is driven by recombined genes that are localized in a couple specific regions of the genome and encode specific functions (selection-driven recombination, and not widespread across the genome). See main text for additional details.

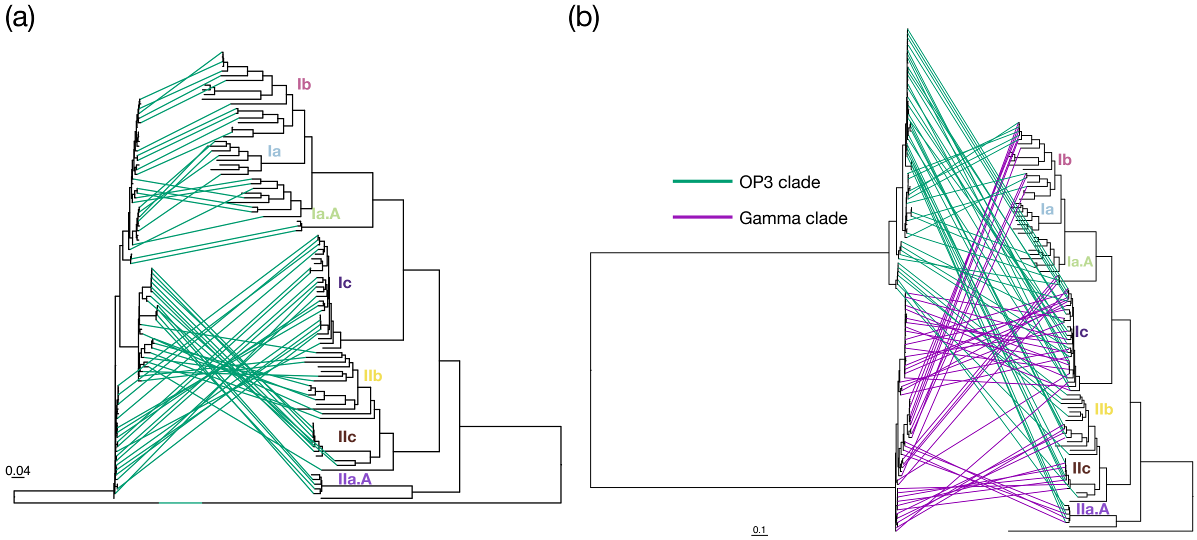


**Figure S4**. **Phylogenetic relationships for all SAGs based on (a) full-length 16S rRNA genes and the whole-genome and (b) NarG protein sequences and the whole genome**. The genome tree was built by extracting, aligning, trimming and concatenating amino acid sequences of universal genes from SAGs and using the resulting alignment with maximum likelihood as implemented in the GToTree software. Scale bar denotes substitution rate.

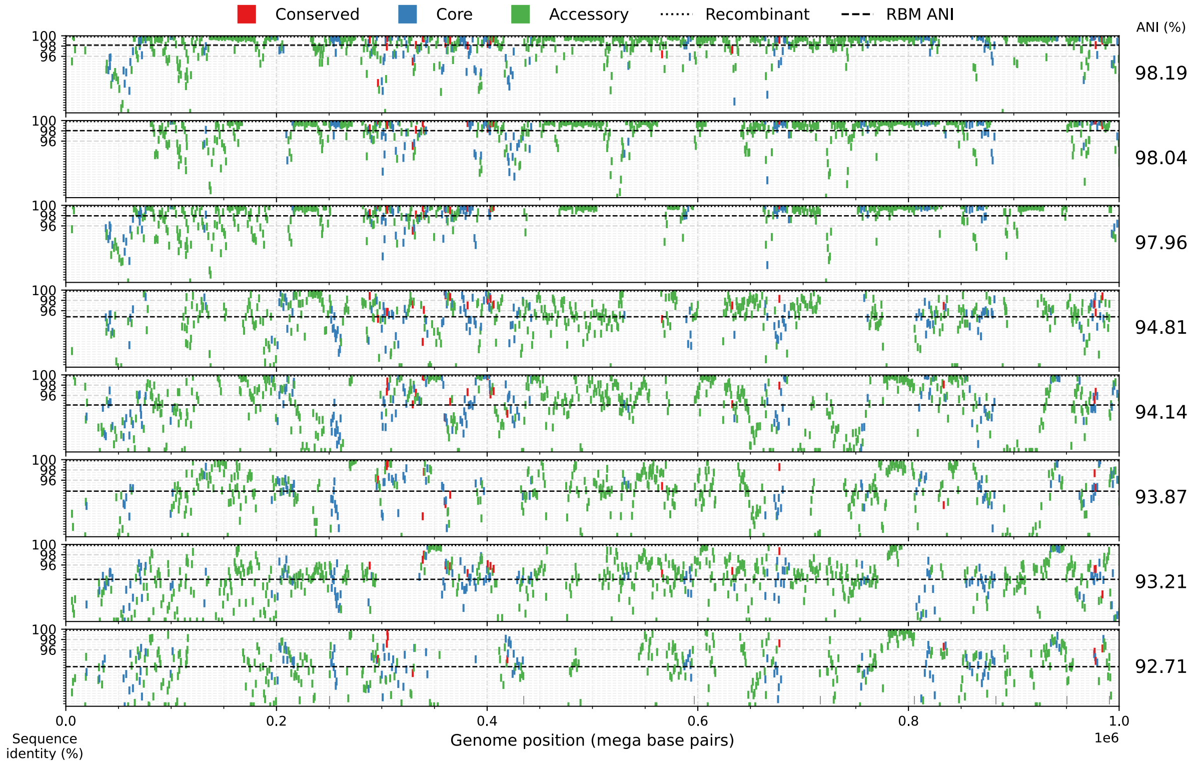


**Figure S5. Extensive recent recombination within the SAR11 subclade Ib**. The figure is identical to Figure 2 except that here eight SAR11 SAGs of the surface subclade Ib were compared against the same (subclade Ib) reference (AG913D08). SAGs were obtained from reference (24). Overall, the picture of subclade Ib is remarkably similar to that of the OMZ subclade Ic (see Figure 2).

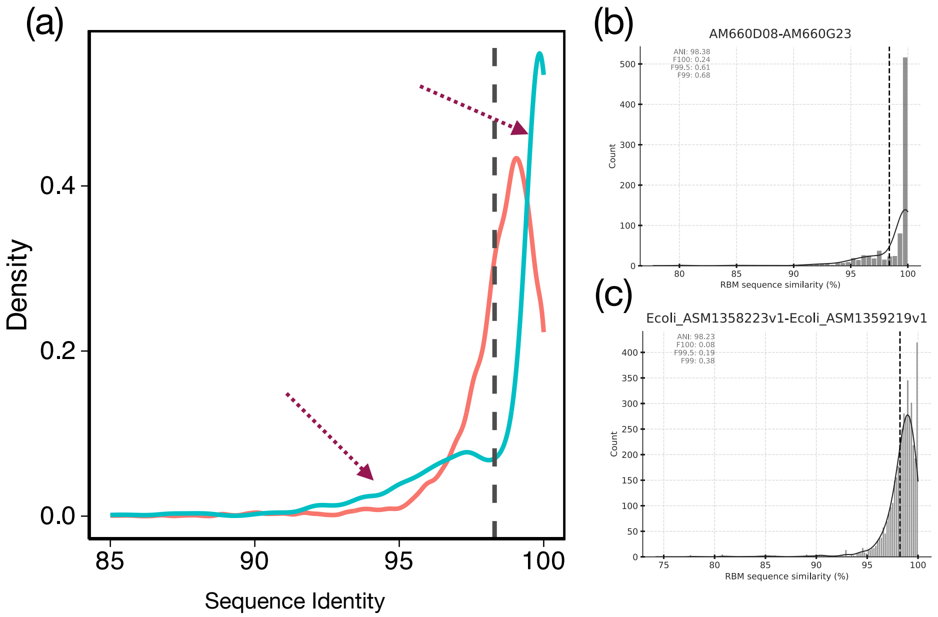


**Figure S6**. **Distribution of nucleotide identities of genes shared between a pair of genomes**. The reciprocal best match genes between two genomes were determined as described in the Methods section and the plots on the right show the number of shared genes (y-axis) plotted against their nucleotide identity (x-axis). The mean of each distribution is essentially the ANI value between the two genomes compared. The plot on the left (**a**) shows the same distribution but the number of genes is expressed as a fraction of total genes shared (normalized to be directly comparable between genomes of different size). The blue line represents a SAR11 SAG subclade Ic genome pair, with its underlying gene distribution shown in (**b**), whereas the red line represents an *E. coli* genome pair, with its underlying gene distribution shown in (**c**); both genome pairs showed about the same ANI, around ~98.3%. Note that the SAR11 SAGs have a significantly higher gene fraction of lower sequence identity shared genes, reflecting horizontal gene transfer mediated by recombination, from more divergent relatives of the species relative to the *E. coli* pair consistent with other results presented elsewhere.

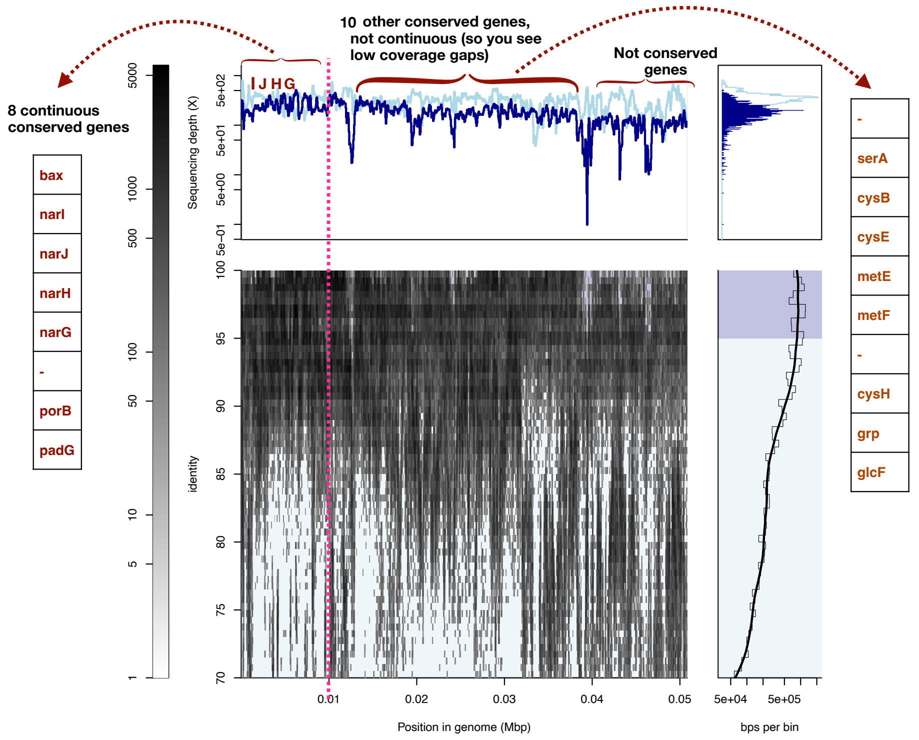


**Figure S7**. **Recruitment plot of the contig containing the *narG* operon**. The figure is identical to Figure S2 except that it shows only the contig that carries the *narG* operon in SAG AM-660-D08 (See Table S1 for information on the corresponding sample where this SAG originated from). This SAG was also used as the reference subclade Ic SAG in Figure 2a. Note the significantly higher identity of reads mapping to the operon relative to those mapping to downstream genes; many of the former reads likely originated from organisms that are not assigned to subclade Ic. Detailed information for the genes carried by the contig and shown on the figure can be found in **Table S3**.

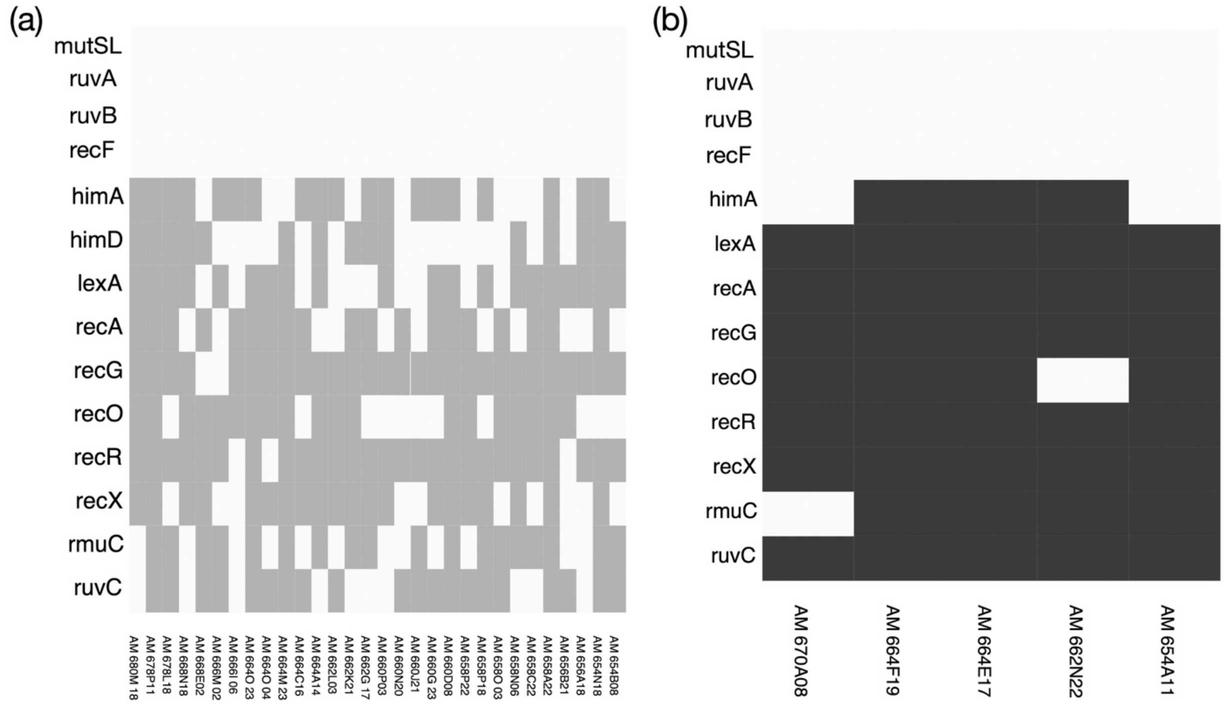


**Figure S8**. **Diversity in gene content related to the process of recombination among the SAG of subclades Ic (a) and IIa.A (b)**. Colored boxes denote that the corresponding gene (row) was found in the genome (column) at a minimum threshold for a match of >35% amino acid identity and >50% alignment ratio. Note that SAGs are incomplete genomes, so the fact that a gene is missing in a couple of the genomes (but is present in several others of the same subclade) is likely due to the incompleteness, although the possibility exists that the SAG truly lacks the corresponding gene. All SAGs seem to be missing the 5 genes shown on the top.

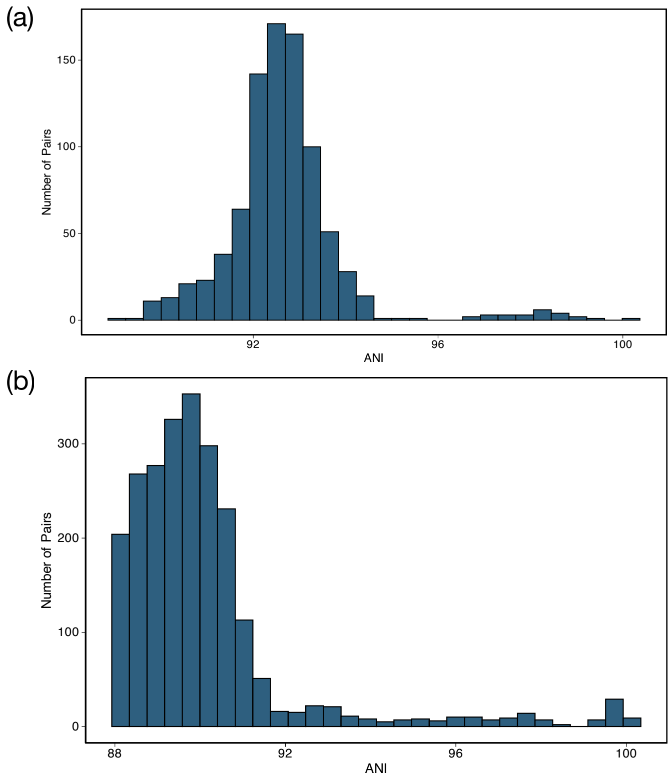


**Figure S9. ANI distribution for SAR11 (a) subclade Ic SAGs and (b) Ib**. A total of 30 SAGs were used that were assigned to subclade Ic (870 comparisons in total). ANI was calculated using FastANI (v1.3.3). The difference with Figure 1b is that it shows all the SAGs from all the clades from the OMZ. Subclade Ib (surface clade) SAGs were obtained from reference (24), a total of 80 SAGs (6320 pairs in total) from the same sampling site (BAT) were used, with genome pairs showing ANI <88% not shown.

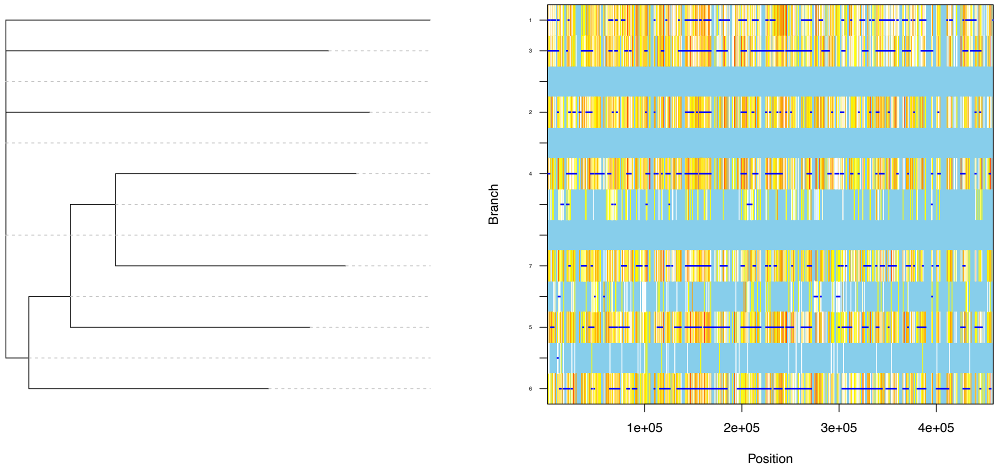


**Figure S10**. **ClonalFrameML analysis of SAR11 subclade Ic SAGs**. Seven SAGs with ANI>95% were used for ClonalFrameML recombination analysis (same genomes as in Figure 2a). Core genome alignment was obtained via ParSNP v1.5.3 and provided to ClonalFrameML. Each vertical line represents a SNP of the alignment, and the horizontal dark blue lines indicate strong signal for recombination for the corresponding genes. Vertical light blue indicates no substitutions; white lines represent no homoplasy while the range from yellow to orange to red reflects increasing degrees of homoplasy. The tree shown was built using the maximum likelihood method as implemented in ClonalFrameML based on the core genome alignment. Average r/m ratios were 3.82 for these SAGs vs. 1.96 for previously published *Salinibacter ruber* and 2.93 for *E. coli* genomes (the same number of genomes, with similar ANI values among the genomes, were chosen for *Salinibacter ruber* and *E. coli*) that were shown to be recombinogenic species (25). It has been shown that ClonalFrameML generally provides similar results with several other methods when r/m ratio is less than 5 (26), as is the case here.

### Supplementary Tables

**Table S1**: Sample collection location, depth and date for the SAG sequences used in this study.

| SAG identifier | Station | Latitude | Longitude | Depth | Collection date |
|---|---|---|---|---|---|
| AM-654 | MEX2 | 18 11 N | 106 17 W | 80 | 3/11/23 |
| AM-656 | MEX2 | 18 11 N | 106 17 W | 95 | 3/6/23 |
| AM-658 | MEX2 | 18 11 N | 106 16 W | 110 | 3/8/23 |
| AM-660 | MEX2 | 18 11 N | 106 17 W | 130 | 3/11/23 |
| AM-662 | MEX2 | 18 11 N | 106 16 W | 140 | 3/8/23 |
| AM-664 | MEX2 | 18 11 N | 106 17 W | 180 | 3/1/23 |
| AM-666 | MEX2 | 18 11 N | 106 16 W | 400 | 3/8/23 |
| AM-668 | MEX2 | 18 10 N | 106 16 W | 600 | 3/4/23 |
| AM-670 | MEX2 | 18 10 N | 106 16 W | 825 | 3/4/23 |
| AM-672 | MEX2 | 18 10 N | 106 16 W | 1000 | 3/4/23 |
| AM-674 | MEX1 | 18 53 N | 104 54 W | 40 | 2/25/23 |
| AM-676 | MEX1 | 18 53 N | 104 54 W | 90 | 2/19/23 |
| AM-678 | MEX1 | 18 53 N | 104 54 W | 120 | 2/17/23 |
| AM-680 | MEX1 | 18 53 N | 104 55 W | 200 | 2/27/23 |

**Table S2**. Clade designation according to the concatenated universal gene tree in Figure 1a and quality of the 105 SAGs used in this study. "Other" denotes that there weren't enough universal genes available that can be extracted from SAGs via the GToTree software for phylogenetic placement. Genome quality was assessed using CheckM v1.5.3 and is reported on columns #3 through #5.

| Genome | Clade | Completeness | Contamination | Strain_heterogeneity |
|---|---|---|---|---|
| AM654A08 | Ia | 71.55 | 0 | 0 |
| AM656A21 | Ia | 52.06 | 1.89 | 100 |
| AM656D15 | Ia | 74.33 | 0.24 | 0 |
| AM674A02 | Ia | 67.25 | 0 | 0 |
| AM674C22 | Ia | 88.73 | 0 | 0 |
| AM674I20 | Ia | 76.12 | 0.47 | 0 |
| AM674J09 | Ia | 75.19 | 0.47 | 0 |
| AM674N09 | Ia | 73.18 | 0 | 0 |
| AM674O03 | Ia | 73.66 | 2.41 | 100 |
| AM674O10 | Ia | 96.21 | 0.47 | 0 |
| AM674O15 | Ia | 93.84 | 0.95 | 0 |
| AM674P21 | Ia | 44.77 | 0 | 0 |
| AM670F15 | Ia.A | 90.77 | 7.23 | 100 |
| AM670L02 | Ia.A | 59.64 | 0 | 0 |
| AM670O04 | Ia.A | 82.37 | 0 | 0 |
| AM670O07 | Ia.A | 52.56 | 0 | 0 |
| AM672C17 | Ia.A | 74.9 | 7.23 | 100 |
| AM672G22 | Ia.A | 62.98 | 0 | 0 |
| AM672I07 | Ia.A | 46 | 0 | 0 |
| AM672M07 | Ia.A | 79.63 | 0 | 0 |
| AM672O05 | Ia.A | 72.55 | 0 | 0 |
| AM654B03 | Ib | 43.98 | 1.2 | 100 |
| AM654C02 | Ib | 74.86 | 0 | 0 |
| AM654D02 | Ib | 70.07 | 0 | 0 |
| AM654G16 | Ib | 65.25 | 0 | 0 |
| AM658A06 | Ib | 44.18 | 1 | 100 |
| AM658N19 | Ib | 46.23 | 0 | 0 |
| AM664C21 | Ib | 71 | 0.71 | 0 |
| AM674M03 | Ib | 54.2 | 0 | 0 |
| AM678O16 | Ib | 58.49 | 0 | 0 |
| AM654B08 | Ic | 61.87 | 0 | 0 |
| AM654N18 | Ic | 74.53 | 0 | 0 |
| AM656A18 | Ic | 54.76 | 0 | 0 |
| AM656B21 | Ic | 59.04 | 0 | 0 |
| AM658A22 | Ic | 93.8 | 0 | 0 |

| | | | | |
|---|---|---|---|---|
| AM658C22 | Ic | 46.49 | 0 | 0 |
| AM658N06 | Ic | 74.36 | 1.2 | 100 |
| AM658O03 | Ic | 81.64 | 0 | 0 |
| AM658P18 | Ic | 68.82 | 0 | 0 |
| AM658P22 | Ic | 49.23 | 0 | 0 |
| AM660D08 | Ic | 82.46 | 0 | 0 |
| AM660G23 | Ic | 80.16 | 0 | 0 |
| AM660N20 | Ic | 55.97 | 0 | 0 |
| AM660P03 | Ic | 70.25 | 0 | 0 |
| AM662G17 | Ic | 59.35 | 0 | 0 |
| AM662K21 | Ic | 72.96 | 1.2 | 100 |
| AM662L03 | Ic | 55.19 | 0 | 0 |
| AM664A14 | Ic | 61.32 | 0 | 0 |
| AM664C16 | Ic | 53.48 | 0 | 0 |
| AM664M23 | Ic | 53 | 0 | 0 |
| AM664O04 | Ic | 55.45 | 0 | 0 |
| AM664O23 | Ic | 79.66 | 0 | 0 |
| AM666I06 | Ic | 49.29 | 0 | 0 |
| AM666M02 | Ic | 64.95 | 0 | 0 |
| AM668E02 | Ic | 84.87 | 0 | 0 |
| AM668N18 | Ic | 93.98 | 0 | 0 |
| AM678F10 | Ic | 60.41 | 0 | 0 |
| AM678L18 | Ic | 90.47 | 0 | 0 |
| AM678P11 | Ic | 85.47 | 0 | 0 |
| AM680M18 | Ic | 59.75 | 0 | 0 |
| AM654A11 | Iia.A | 90.1 | 0 | 0 |
| AM662N22 | IIa.A | 74.92 | 0 | 0 |
| AM664E17 | IIa.A | 87.37 | 1.53 | 25 |
| AM664F19 | IIa.A | 76.94 | 0 | 0 |
| AM670A08 | IIa.A | 87.65 | 0 | 0 |
| AM670C11 | IIb | 94.98 | 0 | 0 |
| AM670C16 | IIb | 94.07 | 0 | 0 |
| AM670D10 | IIb | 56.56 | 0 | 0 |
| AM670E08 | IIb | 78.7 | 0 | 0 |
| AM670E14 | IIb | 97.59 | 0 | 0 |
| AM670G05 | IIb | 71.38 | 0 | 0 |
| AM670O10 | IIb | 53.63 | 0 | 0 |
| AM670P04 | IIb | 71.18 | 0 | 0 |
| AM670P07 | IIb | 72.09 | 1.2 | 100 |
| AM672F18 | IIb | 81.91 | 0 | 0 |
| AM672P15 | IIb | 82.97 | 0 | 0 |
| AM674J13 | IIb | 74.23 | 0 | 0 |
| AM674O23 | IIb | 87.17 | 0 | 0 |
| AM658B22 | IIc | 66.36 | 0 | 0 |
| AM662O06 | IIc | 94.71 | 0 | 0 |
| AM664A04 | IIc | 70.08 | 0 | 0 |
| AM664E23 | IIc | 60.73 | 0 | 0 |
| AM664F23 | IIc | 81.53 | 0 | 0 |
| AM664J17 | IIc | 69.88 | 0 | 0 |
| AM666C10 | IIc | 89.74 | 0 | 0 |
| AM654B15 | Other | 87.95 | 0 | 0 |
| AM654F07 | Other | 87.95 | 0 | 0 |
| AM654K14 | Other | 77.22 | 0 | 0 |
| AM656C17 | Other | 49.95 | 0 | 0 |
| AM656D14 | Other | 91.35 | 0 | 0 |
| AM658C04 | Other | 63.25 | 1.2 | 100 |
| AM658O04 | Other | 77.95 | 0 | 0 |
| AM662K02 | Other | 78.18 | 0 | 0 |
| AM662M16 | Other | 64.94 | 0.11 | 0 |
| AM664D07 | Other | 80.5 | 0 | 0 |
| AM664P01 | Other | 57.76 | 0 | 0 |
| AM666F13 | Other | 58.66 | 0 | 0 |
| AM668B06 | Other | 50.45 | 0 | 0 |
| AM668C15 | Other | 77.11 | 0 | 0 |
| AM670C09 | Other | 74.7 | 0 | 0 |
| AM672N22 | Other | 67.91 | 0 | 0 |
| AM680O15 | Other | 68.13 | 1.2 | 0 |

**Table S3**. Detailed description of the genes found in the recombination hotspot in Figure 2a (2nd red box). Gene annotation was obtained using the eggNOG-mapper v2. Bold fonts denote the *narG* operon.

| Gene ID | pID(nt) | Start | Stop | Strand | COG Category | Gene Annotation | Annotation Description | Alignment Width |
|---|---|---|---|---|---|---|---|---|
| AM660G23_41_7 | 97.587 | 459999 | 460973 | 1 | Conserved Hypothetical | bax | Mannosyl-glycoprotein endo-beta-N-acetylglucosaminidase | 974 |
| **AM660G23_41_6** | **98.406** | **461028** | **461720** | **-1** | **Metabolism 1** | **narI** | **Nitrate reductase gamma subunit** | **692** |

| | | | | | | | | |
|---|---|---|---|---|---|---|---|---|
| **AM660G23_41_5** | **99.149** | **461717** | **462421** | **-1** | **Metabolism 1** | **narJ** | **TIGRFAM nitrate reductase molybdenum cofactor assembly chaperone** | **704** |
| **AM660G23_41_4** | **98.145** | **462418** | **463980** | **-1** | **Metabolism 1** | **narH** | **Respiratory nitrate reductase beta C-terminal** | **1562** |
| **AM660G23_41_3** | **98.81** | **463980** | **467762** | **-1** | **Metabolism 1** | **narG** | **Respiratory nitrate reductase alpha N-terminal** | **3782** |
| AM662K21_74_2 | 99.763 | 467791 | 469479 | -1 | Metabolism 1 | - | NADPH-dependent glutamate synthase beta | 1688 |
| AM660N20_12_10 | 99.906 | 469479 | 470543 | -1 | Metabolism 1 | porB | Thiamine pyrophosphate enzyme, C-terminal TPP binding domain | 1064 |
| AM660N20_12_13 | 99.32 | 472635 | 473516 | -1 | Metabolism 1 | - | phosphoserine phosphatase | 881 |
| AM660N20_12_14 | 98.169 | 473506 | 475089 | -1 | Metabolism 1 | serA | D-isomer specific 2-hydroxyacid dehydrogenase, catalytic domain | 1583 |
| AM662K21_34_7 | 97.305 | 476251 | 477252 | -1 | Metabolism 1 | cysB | Pyridoxal-phosphate dependent enzyme | 1001 |
| AM662L03_33_7 | 97.631 | 477283 | 477873 | -1 | Metabolism 1 | cysE | TIGRFAM serine O-acetyltransferase | 590 |
| AM660N20_12_21 | 98.596 | 480480 | 482759 | -1 | Metabolism 1 | metE | Cobalamin-independent synthase, Catalytic domain | 2279 |
| AM660N20_12_22 | 98.105 | 482769 | 483665 | -1 | Metabolism 1 | metF | Methylenetetrahydrofolate reductase | 896 |
| AM662L03_14_23 | 98.438 | 484221 | 484739 | -1 | Conserved Hypothetical | - | Protein conserved in bacteria | 518 |
| AM662K21_38_4 | 99.602 | 488277 | 489029 | -1 | Metabolism 1 | cysH | Belongs to the PAPS reductase family. CysH subfamily | 752 |
| AM664M23_15_8 | 98.475 | 489351 | 489809 | 1 | Information | grp | helix_turn_helix ASNC type | 458 |
| AM662L03_14_15 | 98.225 | 491656 | 492951 | -1 | Metabolism 1 | glcF | 4Fe-4S dicluster domain | 1295 |

- means no gene annotation was provided from the EggNog database.

**Table S4.** Detailed description of the genes found in the recombination hotspot in Figure 2a (3rd red box). Gene annotation was obtained using the eggNOG-mapper v2.

| Gene ID | pID(nt) | Start | Stop | Strand | COG Category | Gene Annotation | Annotation Description | Alignment Width |
|---|---|---|---|---|---|---|---|---|
| AM660G23_36_5 | 100 | 907436 | 907621 | 1 | Hypothetical | - | Hypothetical | 185 |
| AM660G23_36_7 | 99.815 | 908243 | 909322 | 1 | Metabolism 2 | - | Part of the tripartite ATP-independent periplasmic (TRAP) transport system | 1079 |
| AM660G23_36_9 | 100 | 910163 | 910252 | 1 | Hypothetical | - | Hypothetical | 89 |
| AM660J21_5_6 | 100 | 910252 | 910977 | 1 | Metabolism 1 | phbB | KR domain | 725 |
| AM660N20_19_5 | 99.248 | 911001 | 912197 | 1 | Metabolism 1 | phbA | Belongs to the thiolase family | 1196 |
| AM660J21_5_9 | 98.485 | 914119 | 915108 | 1 | Metabolism 1 | ldhA | PFAM D-isomer specific 2-hydroxyacid dehydrogenase, catalytic region | 989 |

**Table S5.** Sample information for the metagenomes from the ETNP OMZ used in this study.

| Sample ID | DNA Concentration(µm/L) | Date Sampled | Date Extracted | Lat N | Long W | Depth (m) | Volume Filtered (mL) | Station |
|---|---|---|---|---|---|---|---|---|
| 1053 | 6.92 | 12/30/2021 | 10.16.22 | 12º45.662N | 90º26.980W | 601.1 | 2250 | 20 |
| 1055 | 6.16 | 12/30/2021 | 9.23.22 | 12º45.662N | 90º26.980W | 501.2 | 2200 | 20 |
| 1056 | 3.24 | 12/30/2021 | 10.2.22 | 12º45.662N | 90º26.980W | 451.3 | 2020 | 20 |
| 1059 | 3.32 | 12/30/2021 | 10.2.22 | 12º45.662N | 90º26.980W | 401.5 | 1950 | 20 |
| 1061 | 5.6 | 12/30/2021 | 9.23.22 | 12º45.662N | 90º26.980W | 351.5 | 2250 | 20 |
| 1063 | 7.32 | 12/30/2021 | 9.30.22 | 12º45.662N | 90º26.980W | 331.3 | 2200 | 20 |
| 1065 | 12 | 12/30/2021 | 10.12.22 | 12º45.662N | 90º26.980W | 321.6 | 2250 | 20 |
| 1067 | 4.52 | 12/30/2021 | 10.2.22 | 12º45.662N | 90º26.980W | 306.7 | 2250 | 20 |
| 1069 | 3.82 | 12/30/2021 | 9.30.22 | 12º45.662N | 90º26.980W | 251.9 | 2300 | 20 |
| 1088 | 4.24 | 01/02/2022 | 10.12.22 | 11º48.729N | 94º20.501W | 551.8 | 2100 | 28 |
| 1090 | 6.36 | 01/02/2022 | 10.12.22 | 11º48.729N | 94º20.501W | 451.9 | 2050 | 28 |
| 1092 | 8.72 | 01/02/2022 | 9.30.22 | 11º48.729N | 94º20.501W | 372 | 2150 | 28 |
| 1094 | 8.8 | 01/02/2022 | 10.16.22 | 11º48.729N | 94º20.501W | 302.1 | 2000 | 28 |
| 1096 | 8.36 | 01/02/2022 | 10.2.22 | 11º48.729N | 94º20.501W | 273 | 2160 | 28 |
| 1098 | 6.12 | 01/02/2022 | 10.2.22 | 11º48.729N | 94º20.501W | 253 | 2240 | 28 |
| 1100 | 5.4 | 01/02/2022 | 9.30.22 | 11º48.729N | 94º20.501W | 227 | 2300 | 28 |

| | | | | | | | | |
|---|---|---|---|---|---|---|---|---|
| 1102 | 12.8 | 01/02/2022 | 10.12.22 | 11º48.729N | 94º20.501W | 173 | 2200 | 28 |
| 1104 | 8.32 | 01/02/2022 | 10.16.22 | 11º48.729N | 94º20.501W | 113 | 2160 | 28 |
| 1144 | 2.3 | 01/08/2022 | 10.2.22 | 12º47.379N | 102º25.779W | 502 | 2250 | 38 |
| 1146 | 7.44 | 01/08/2022 | 9.30.22 | 12º47.379N | 102º25.779W | 401 | 2060 | 38 |
| 1148 | 8.76 | 01/08/2022 | 10.12.22 | 12º47.379N | 102º25.779W | 352 | 2150 | 38 |
| 1150 | 11 | 01/08/2022 | 10.16.22 | 12º47.379N | 102º25.779W | 302 | 2100 | 38 |
| 1157 | 13 | 01/08/2022 | 9.30.22 | 12º47.379N | 102º25.779W | 282 | 2400 | 38 |
| 1159 | 4.84 | 01/08/2022 | 10.12.22 | 12º47.379N | 102º25.779W | 267 | 2180 | 38 |
| 1152 | 3.35 | 01/08/2022 | 9.23.22 | 12º47.379N | 102º25.779W | 202 | 2260 | 38 |
| 1154 | 5.6 | 01/08/2022 | 10.2.22 | 12º47.379N | 102º25.779W | 183 | 2340 | 38 |
| 1160 | 8.48 | 01/08/2022 | 9.30.22 | 12º47.379N | 102º25.779W | 133 | 2230 | 38 |
| 1170 | 4.16 | 01/10/2022 | 9.23.22 | 13º39.016N | 103º59.970W | 501 | 2240 | 40 |
| 1172 | 5.6 | 01/10/2022 | 9.30.22 | 13º39.016N | 103º59.970W | 402 | 2000 | 40 |
| 1174 | 3.98 | 01/10/2022 | 10.16.22 | 13º39.016N | 103º59.970W | 351 | 2220 | 40 |
| 1176 | 10 | 01/10/2022 | 10.12.22 | 13º39.016N | 103º59.970W | 301 | 2250 | 40 |
| 1178 | 12.3 | 01/10/2022 | 10.2.22 | 13º39.016N | 103º59.970W | 286 | 2380 | 40 |
| 1180 | 5.08 | 01/10/2022 | 10.2.22 | 13º39.016N | 103º59.970W | 272 | 2180 | 40 |
| 1181 | 4.68 | 01/10/2022 | 9.30.22 | 13º39.016N | 103º59.970W | 202 | 2220 | 40 |
| 1184 | 7.44 | 01/10/2022 | 10.12.22 | 13º39.016N | 103º59.970W | 147 | 2320 | 40 |
| 1186 | 9.4 | 01/10/2022 | 9.23.22 | 13º39.016N | 103º59.970W | 122 | 2600 | 40 |
| 1218 | 3.69 | 01/14/2022 | 9.30.22 | 16º58.790N | 107º42.002W | 402 | 2150 | 45 |
| 1220 | 4.52 | 01/14/2022 | 9.23.22 | 16º58.790N | 107º42.002W | 302 | 2120 | 45 |
| 1222 | 6.92 | 01/14/2022 | 10.2.22 | 16º58.790N | 107º42.002W | 252 | 2040 | 45 |
| 1224 | 6.8 | 01/14/2022 | 10.12.22 | 16º58.790N | 107º42.002W | 202 | 2120 | 45 |
| 1237 | 6.72 | 01/14/2022 | 10.12.22 | 16º58.790N | 107º42.002W | 202 | 4060 | 45 |
| 1226 | 5.76 | 01/14/2022 | 10.2.22 | 16º58.790N | 107º42.002W | 177 | 2360 | 45 |
| 1228 | 10.5 | 01/14/2022 | 10.12.22 | 16º58.790N | 107º42.002W | 153 | 2150 | 45 |
| 1238 | 17.3 | 01/14/2022 | 10.16.22 | 16º58.790N | 107º42.002W | 153 | 4060 | 45 |
| 1230 | 7.48 | 01/14/2022 | 9.23.22 | 16º58.790N | 107º42.002W | 132 | 1820 | 45 |
| 1239 | 20.5 | 01/14/2022 | 10.16.22 | 16º58.790N | 107º42.002W | 132 | 4380 | 45 |
| 1231 | 7.88 | 01/14/2022 | 9.30.22 | 16º58.790N | 107º42.002W | 122 | 1900 | 45 |
| 1234 | 6.48 | 01/14/2022 | 10.16.22 | 16º58.790N | 107º42.002W | 112 | 2180 | 45 |

**References cited**.


1. D. Tsementzi *et al.*, SAR11 bacteria linked to ocean anoxia and nitrogen loss. *Nature* **536**, 179-183 (2016).

2. M. R. Lindsay *et al.*, Species-resolved, single-cell respiration rates reveal dominance of sulfate reduction in a deep continental subsurface ecosystem. *Proceedings of the National Academy of Sciences* **121**, e2309636121 (2024).

3. A. Bankevich *et al.*, SPAdes: a new genome assembly algorithm and its applications to single-cell sequencing. *Journal of computational biology* **19**, 455-477 (2012).

4. R. Stepanauskas *et al.*, Improved genome recovery and integrated cell-size analyses of individual uncultured microbial cells and viral particles. *Nature Communications* **8**, 84 (2017).

5. J. Zhao *et al.*, Microbial Response to Natural Disturbances: Rare Biosphere often plays a role. *bioRxiv* (2024).

6. L. M. Rodriguez-R *et al.*, The Microbial Genomes Atlas (MiGA) webserver: taxonomic and gene diversity analysis of Archaea and Bacteria at the whole genome level. *Nucleic Acids Research* **46**, W282-W288 (2018).

7. L. M. Rodriguez-r, K. T. Konstantinidis, Nonpareil: a redundancy-based approach to assess the level of coverage in metagenomic datasets. *Bioinformatics* **30**, 629-635 (2014).

8. Y. Peng, H. C. Leung, S.-M. Yiu, F. Y. Chin, IDBA-UD: a de novo assembler for single-cell and metagenomic sequencing data with highly uneven depth. *Bioinformatics* **28**, 1420-1428 (2012).

9. L. M. Rodriguez-R, K. T. Konstantinidis (2016) The enveomics collection: a toolbox for specialized analyses of microbial genomes and metagenomes. (PeerJ Preprints).

10. G. V. Uritskiy, J. DiRuggiero, J. Taylor, MetaWRAP—a flexible pipeline for genome-resolved metagenomic data analysis. *Microbiome* **6**, 1-13 (2018).

11. C. M. Sieber *et al.*, Recovery of genomes from metagenomes via a dereplication, aggregation and scoring strategy. *Nature microbiology* **3**, 836-843 (2018).

12. D. H. Parks, M. Imelfort, C. T. Skennerton, P. Hugenholtz, G. W. Tyson, CheckM: assessing the quality of microbial genomes recovered from isolates, single cells, and metagenomes. *Genome research* **25**, 1043-1055 (2015).

13. J. Zhao, J. P. Both, L. M. Rodriguez-R, K. T. Konstantinidis, GSearch: ultra-fast and scalable genome search by combining K-mer hashing with hierarchical navigable small world graphs. *Nucleic Acids Research*, gkae609 (2024).

14. D. H. Parks *et al.*, GTDB: an ongoing census of bacterial and archaeal diversity through a phylogenetically consistent, rank normalized and complete genome-based taxonomy. *Nucleic acids research* **50**, D785-D794 (2022).

15. S. F. Altschul, W. Gish, W. Miller, E. W. Myers, D. J. Lipman, Basic local alignment search tool. *Journal of molecular biology* **215**, 403-410 (1990).

16. X. Didelot, D. J. Wilson, ClonalFrameML: efficient inference of recombination in whole bacterial genomes. *PLoS computational biology* **11**, e1004041 (2015).

17. M. D. Lee, GToTree: a user-friendly workflow for phylogenomics. *Bioinformatics* **35**, 4162-4164 (2019).

18. C. Jain, R. L. Rodriguez, A. M. Phillippy, K. T. Konstantinidis, S. Aluru, High throughput ANI analysis of 90K prokaryotic genomes reveals clear species boundaries. *Nat Commun* **9**, 5114 (2018).

19. B. Q. Minh *et al.*, IQ-TREE 2: new models and efficient methods for phylogenetic inference in the genomic era. *Molecular biology and evolution* **37**, 1530-1534 (2020).

20. R. C. Edgar, MUSCLE: multiple sequence alignment with high accuracy and high throughput. *Nucleic acids research* **32**, 1792-1797 (2004).

21. C. P. Cantalapiedra, A. Hernández-Plaza, I. Letunic, P. Bork, J. Huerta-Cepas, eggNOG-mapper v2: functional annotation, orthology assignments, and domain prediction at the metagenomic scale. *Molecular biology and evolution* **38**, 5825-5829 (2021).

22. A. Hernández-Plaza *et al.*, eggNOG 6.0: enabling comparative genomics across 12 535 organisms. *Nucleic Acids Research* **51**, D389-D394 (2023).

23. L.-M. Rodriguez-R, and Konstantinidis, K. T., The enveomics collection: a toolbox for specialized analyses of microbial genomes and metagenomes. *PeerJ Preprints* (2016).

24. M. G. Pachiadaki *et al.*, Charting the complexity of the marine microbiome through single-cell genomics. *Cell* **179**, 1623-1635. e1611 (2019).

25. R. E. Conrad, Brink, C. E., Viver, T., Rodriguez-R, L. M., Aldeguer-Riquelme, B., Hatt, J. K., Venter, S. N., Rossello-Mora, R., Amann, R., and Konstantinidis, K. T., Microbial species and intraspecies units exist 1 and are maintained by ecological cohesivenesscoupled to high homologous recombination. *Nature Communications*, In press. (2024).

26. E. L. Torrance, C. Burton, A. Diop, L.-M. Bobay, Evolution of homologous recombination rates across bacteria. *Proceedings of the National Academy of Sciences* **121**, e2316302121 (2024).
